# Sex-dependent changes in murine striatal dopamine release, sleep, and behavior during spontaneous Δ-9-tetrahydrocannabinol abstinence

**DOI:** 10.1101/2021.12.01.470765

**Authors:** Andrew J. Kesner, Yolanda Mateo, Karina P. Abrahao, Stephanie Ramos-Maciel, Matthew J. Pava, Alexa L. Gracias, Riley T. Paulsen, Hartley B. Carlson, David M. Lovinger

## Abstract

Withdrawal symptoms are observed upon cessation of cannabis use in humans. Although animal studies have examined withdrawal symptoms following exposure to delta-9-tetrahydrocannabinol (THC), difficulties in obtaining objective measures of spontaneous withdrawal using paradigms that mimic cessation of use in humans have slowed research. The neuromodulator dopamine (DA) is known to be affected by chronic THC treatment and plays a role in many behaviors related to human THC withdrawal symptoms. These symptoms include sleep disturbances that often drive relapse, and emotional behaviors, e.g., irritability and anhedonia. We examined THC withdrawal-induced changes in striatal DA release and the extent to which sleep disruption and behavioral maladaptation manifest during withdrawal in a mouse chronic cannabis exposure model. Using a THC treatment regimen known to produce tolerance we measured electrically elicited DA release in acute brain slices from different striatal subregions during early and late THC abstinence. Long-term polysomnographic recordings from mice were used to assess vigilance state and sleep architecture before, during, and after THC treatment. We additionally assessed how behaviors that model human withdrawal symptoms are altered by chronic THC treatment in early and late abstinence. We detected altered striatal DA release, sleep disturbances that mimic clinical observations, and behavioral maladaptation in mice following tolerance inducing THC treatment. Sex differences were observed in nearly all metrics. Altered striatal DA release, sleep and affect-related behaviors associated with spontaneous THC abstinence were more consistently observed in male mice. To our knowledge these findings provide the first model of directly translatable non-precipitated cannabis withdrawal symptoms, in particular, sleep disruption.

## Introduction

### Cannabis use and abuse

Cannabis derivatives are currently the most widely used illicit drugs in the world [1], with reported use increasing, at least in part due to efforts towards its legalization [2]. While adverse side-effects of cannabis use are often mild, a subpopulation of chronic cannabis users experience hallmarks of a substance use disorder (SUD), including use despite adverse consequences, craving, tolerance to the drug’s effects, and withdrawal symptoms [3–5]. Experiencing these symptoms can lead to diagnosis of cannabis use disorder (CUD) as defined by the DSM-5 [6].

The main psychoactive component of cannabis is delta-9-tetrahydrocannabinol (THC). This compound is primarily responsible for the drug-induced “high” via actions at the cannabinoid-type 1 receptor (CB1) [7, 8]. Indeed, behavioral effects of chronic THC treatment are absent in mice genetically engineered to lack the CB1 receptor [9, 10], and CB1 antagonists prevent self-administration of synthetic cannabinoids [10, 11].

### Cannabis withdrawal symptoms

Cessation of chronic cannabis or THC use causes withdrawal symptoms in a significant population of users [12–25]. In humans, cannabis withdrawal symptoms (CWS) may include: irritability/aggression, nervousness/anxiety, disrupted sleep, hypophagia and weight loss, restlessness, depressed mood, uncomfortable somatic symptoms e.g. abdominal pain, shakes, sweating, fever/chills, and headache [6]. One of the most prominent CWS is disrupted sleep, and poor sleep quality is a major risk factor towards cannabis relapse [26–28]. Human studies show that acute THC produces sleep facilitation including shorter sleep latency, less difficulty falling asleep and more time spent in sleep [29–32]. Our group has shown that endocannabinoid (eCB) activity contributes to non-rapid eye movement, (NREM) stability [33]. The sleep-inducing effects of acute THC may dissipate during chronic exposure in humans [29], and withdrawal from chronic THC or cannabis use in humans is associated with a decrease in sleep time and efficiency [34–36].

At the preclinical level, CWS have been modeled using spontaneous withdrawal (abrupt cessation of drug treatment). However, withdrawal precipitated by CB1 antagonist administration, which immediately blocks the effects of the chronically administered cannabinoid drug, is the more prevalent model [37]. An obvious caveat of precipitated withdrawal is that humans undergo spontaneous withdrawal, so the translational relevance of precipitated CWS is unclear. Spontaneous cannabis or THC withdrawal reliably induces somatic withdrawal symptoms (e.g. wet dog and head shaking, front paw tremor, hunched posture, body tremor, etc.) [38–41], yet previous studies have struggled to provide strong evidence for spontaneous THC withdrawal symptoms that more closely model human CWS.

### Dopamine, CUD and CWS

The neuromodulator dopamine (DA) has well established roles in SUDs [42, 43], behavioral alterations during drug withdrawal [44, 45], and sleep [46–48].There is considerable evidence that exogenous cannabinoids indirectly modulate DA activity [49–51] with ramifications for development of CUD and CWS. The striatum, comprised of subregions including the nucleus accumbens (NAc), dorsal medial (DMS) and dorsal lateral striatum (DLS), is a major site of DA action in the brain and is involved in many behaviors associated with drug abuse and withdrawal symptoms [52], as well as sleep [53], making it an interesting nexus of the actions of THC and withdrawal phenomena.

### Aim of study

In the present study, we modeled chronic cannabis use using a well-established procedure to induce THC tolerance in mice [54] to examine whether spontaneous THC withdrawal drives: 1) alterations in striatal DA release, 2) sleep disruption, and 3) translationally relevant behavioral maladaptation. To examine effects of acute and chronic THC treatment on behavior and sleep we compared vehicle injections to the first and last injections of the chronic THC or vehicle control (VEH) treatment. To examine maladaptive behavior and sleep during withdrawal we compared changes in baseline metrics (2 days pretreatment) to early (1-2 days) and late (5-6 days) of abstinence in mice treated with THC or VEH (**Supplemental Figure 1**). Both male and female mice were examined, as accumulating evidence suggests sex-differences exist in the development of CUD and severity of CWS [55].

## Methods and Materials

### Subjects

All experiments were conducted using wildtype, 8 to 10-week-old male and female mice (Mus musculus, C57BL/6J strain) at the time of electroencephalogram/electromyogram (EEG/EMG) implantation or behavioral studies. For full description of subjects see **supplemental information**. All methods used in this work were approved by the Animal Care and Use Committee of the National Institute on Alcohol Abuse and Alcoholism (protocol #: LIN-DL-1) and were within the guidelines described in the NIH Guide to the Care and Use of Laboratory Animals.

### Sleep recordings

Sleep experiments were performed as previously described [33, 56]. See **supplemental information** for full description of surgical procedures, recording environment, polysomnographic acquisition, and vigilance-state scoring (e.g., NREM vs REM sleep).

### THC treatment and behavioral paradigms

#### Sleep effects during and after THC treatment

Recordings and treatment began after the 7-day acclimatization period. First, we collected 2 separate days of baseline polysomnographic measurements (pretreatment). Following stoppage of recording on the second day of baseline, all mice received injection of VEH consisting of 1:1:18 (Dimethysulfoxide (DMSO) : Cremaphor : 0.9% Saline) (Cat# D2650 and C5135 for DMSO and Cremaphor EL, respectively, Sigma Aldrich, St. Louis, MO). After stoppage of recording for this VEH treatment, mice were treated with either VEH or 10mg/kg of THC in vehicle and EEG/EMG were recorded for another ∼22 hrs. After stoppage of recording for first injection the animals received injections according to their assigned group twice daily for four more days – once before the dark phase, between ZT11:00 and ZT12:00, and the second injection between ZT1:00 and ZT2:00. Treatments were assigned randomly such that each recording chamber, which housed five individually housed mice, had roughly half the subjects receiving either THC or VEH. Injections for Vehicle-only session and first injection occurred within the last hour before the start of the dark phase, between zeitgeber time (i.e. lights on; ZT) ZT11:00 and ZT12:00. Finally, the mice received one last injection, again between ZT11:00 and ZT12:00, and EEG/EMG were recorded for ∼22 hr. We then continued recording EEG/EMG for the next six full days of abstinence (ABST 1-6). Each ABST recording began between ZT11:00 and ZT12:00, and lasted ∼22 hrs. To measure effects from chronic low dose THC, the same procedure as above was used, but a 1mg/kg dose of THC was given. To measure sleep effects during abstinence from an acute dose of THC, the same procedure as chronic 10 mg/kg THC experiments was used except a single THC treatment was immediately followed by ABST 1-6.

#### Behavioral effects of chronic 10 mg/kg dose of THC

For all awake behaviors, mice were treated with the chronic 10 mg/kg dose of THC using the treatment procedure described above in sleep experiments. All mice were initially group housed, 4 animals per cage. Before all experiments mice were transported to respective behavioral rooms and handled daily for 3 days.

See **supplemental information** for full description of sucrose preference test, operant conditioned sucrose seeking, intake and locomotor activity metrics, and bottle brush test procedures.

### Plasma corticosterone quantification

Blood was collected via lateral tail vein bleeding. See **supplemental information** for full description of plasma corticosterone collection procedures. Plasma corticosterone concentration was determined using enzyme-linked immunosorbent assay as per manufacturer’s instructions for small sample volume (ELISA part # ADI-901-097, Enzo Life Sciences, Inc., NY). The assay was performed in duplicate and read using a SpectraMax190 Microplate Reader. Standard curves and corticosterone concentration extrapolation were performed using an open-access online analysis service (MyAssays Ltd., USA).

### Statistical analysis

Data were analyzed using GraphPad Prism (version 9; GraphPad Software, La Jolla, CA, USA) statistics software. All behavioral data and CORT measurements were transformed to change, “Δ”, from vehicle injection (when investigating effects of first and last treatments) or pretreatment (when investigating effects during early and late abstinence). See **supplemental information** for full description of statistical analysis.

## Results

### Striatal dopamine release in early and late abstinence after acute and chronic THC treatment

#### Sex-dependent changes in DA release during abstinence following acute THC treatment

We first measured electrically stimulated DA release in the NAc, DMS, and DLS during early and late abstinence following acute THC treatment. Male mice showed a significant increase in peak DA release in DMS following a single, i.e., acute, injection of THC (**Figure 1A**). This significant increase was observed during early (**Figure 1B**) and late abstinence (**Figure 1C**). However, there were no differences in peak DA release in the DLS or NAc after an acute THC injection. In contrast, female mice showed decreased peak DA levels in DMS and NAc following a single THC dose in early abstinence. However, this same THC administration protocol elicited an increase in peak DA levels in DLS (**Figure 1D**). In late abstinence after acute THC treatment no differences in peak DA release were observed (**Figure 1E**). THC treatments did not elicit changes in uptake rate (data not shown).

**Figure 1.**
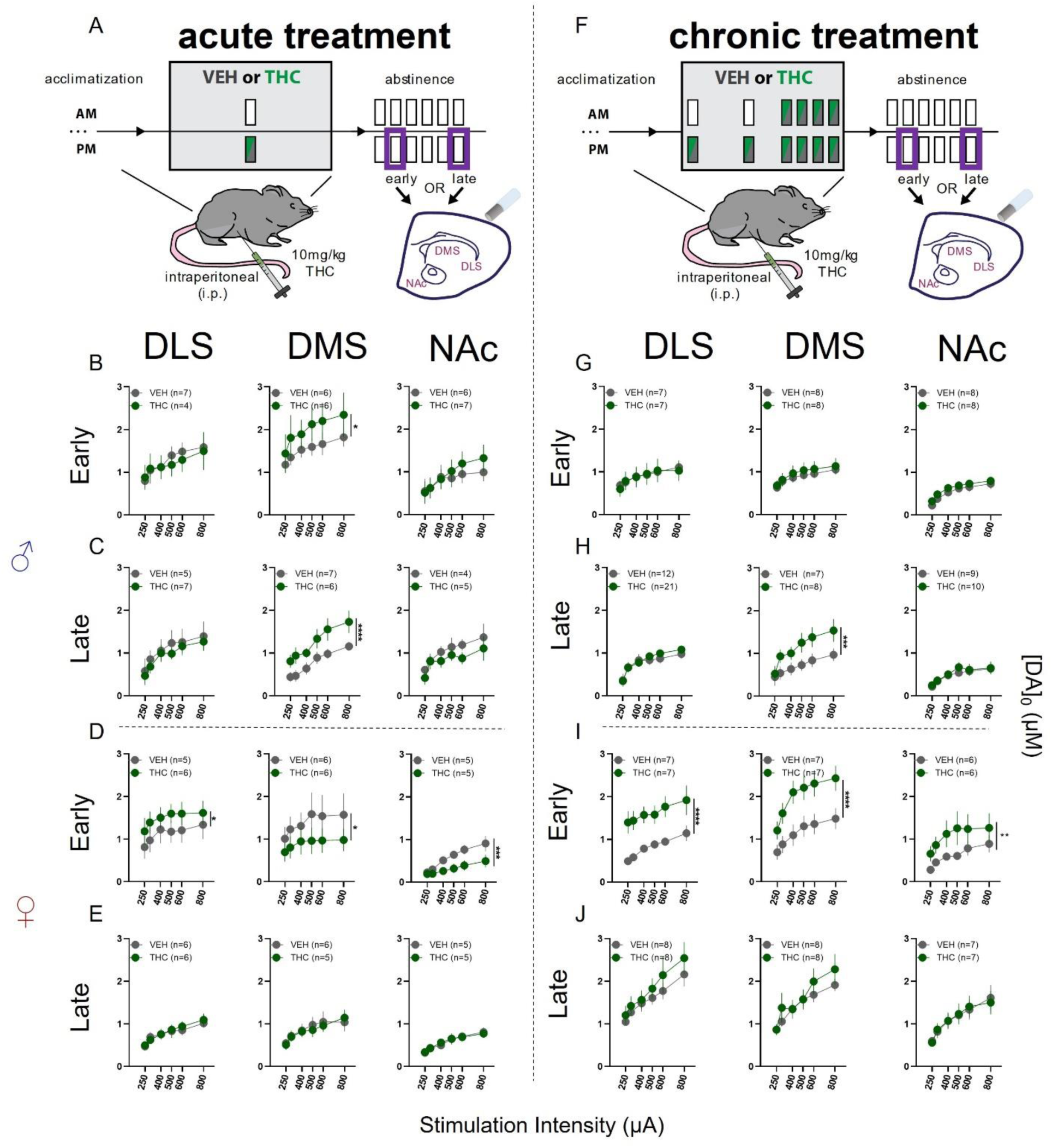
Effects of acute or chronic THC treatment on striatal DA release in acute brain slices using FSCV. (A) Schematic illustrating acute treatment and time points of FSCV recordings in striatal subregions. (B) DA release in males during early abstinence following acute treatment. No differences detected in DLS or NAc. In DMS THC treated mice had elevated DA release (F_treatment_(1,60) = 4.866; p = 0.0312) (C) DA release in males during late abstinence following acute treatment. No differences detected in DLS or NAc. In DMS THC treated mice had elevated DA release (F_treatment_(1,59) = 21.33; p = <0.0001) (D) DA release in females during early abstinence following acute treatment. In DLS THC treated mice had elevated DA release (F_treatment_(1,54) = 4.726, p = 0.0341), while DA release was reduced in DMS (F_treatment_ (1,60) = 5.669; p = 0.02) and NAc (F_treatment_(1,48) = 15.87; p = 0.0002). (E) DA release in females during late abstinence following acute treatment. No differences detected in DLS, DMS, or NAc. (F) Schematic illustrating chronic treatment and time points of FSCV recordings in striatal subregions. (G) DA release in males during early abstinence following chronic treatment. No differences detected in DLS, DMS, or NAc. (H) DA release in males during late abstinence following chronic treatment. No differences detected in DLS or NAc. In DMS THC treated mice had elevated DA release (F_treatment_ (1,71) = 13.56; p = 0.0004). (I) DA release in females during early abstinence following chronic treatment. DA release was elevated in DLS (F_treatment_(1,72) = 51.61, p = ˂0.0001), DMS (F_treatment_(1,72) = 33.82, p = ˂0.0001) and NAc (F_treatment_(1,60) = 11.69, p = 0.0011). (J) DA release in females during late abstinence following acute treatment. No differences detected in DLS, DMS, or NAc.

#### Sex-dependent changes in DA release during abstinence following chronic THC treatment

In male mice exposed to chronic THC treatment (**Figure 1F**), a significant increase in DMS DA release was present during late abstinence but not early abstinence (**Figure 1G-H**). In females, during early abstinence we found a robust increase in DA levels in all three striatal regions (**Figure 1I**), but no such increase during late abstinence (**Figure 1J**).

### Sleep and vigilance state architecture during acute and chronic THC treatment and abstinence

We next examined if THC and VEH treatments changed percent time spent in NREM and REM sleep and sleep architecture (i.e., bout number and duration) referenced to baseline metrics (i.e., vehicle treatment or pretreatment epochs). Note that the sleep and behavioral data that follow are reported as a change-score (i.e. delta “Δ”) from either Vehicle-treatment or pretreatment epochs.

#### Acute and chronic THC exposure alters sleep in a sex dependent manner

We first examined how VEH or THC treatments altered sleep after the first and last injections of the chronic treatment regimen as compared to the vehicle treatment session (**Figure 2A and B**). In male mice, the first THC injection increased percent time spent in NREM compared to VEH during the dark photoperiod (lights off; LOFF). After the last injection, percent time in NREM was lower in THC treated compared to VEH treated male mice during both LOFF and the light photoperiod (lights on; LON). During LOFF, THC-treated male mice showed a lesser change in percent time in NREM as compared to the first injection (**Figure 2C**). No significant effects were detected for NREM bout duration (**Figure 2D**). However, an increased number of NREM bouts was observed during LOFF in THC-treated male mice compared to VEH-treated mice. After the last injection the number of NREM bouts during both LOFF and LON was decreased regardless of treatment when compared to the first injection (**Figure 2E**).

**Figure 2.**
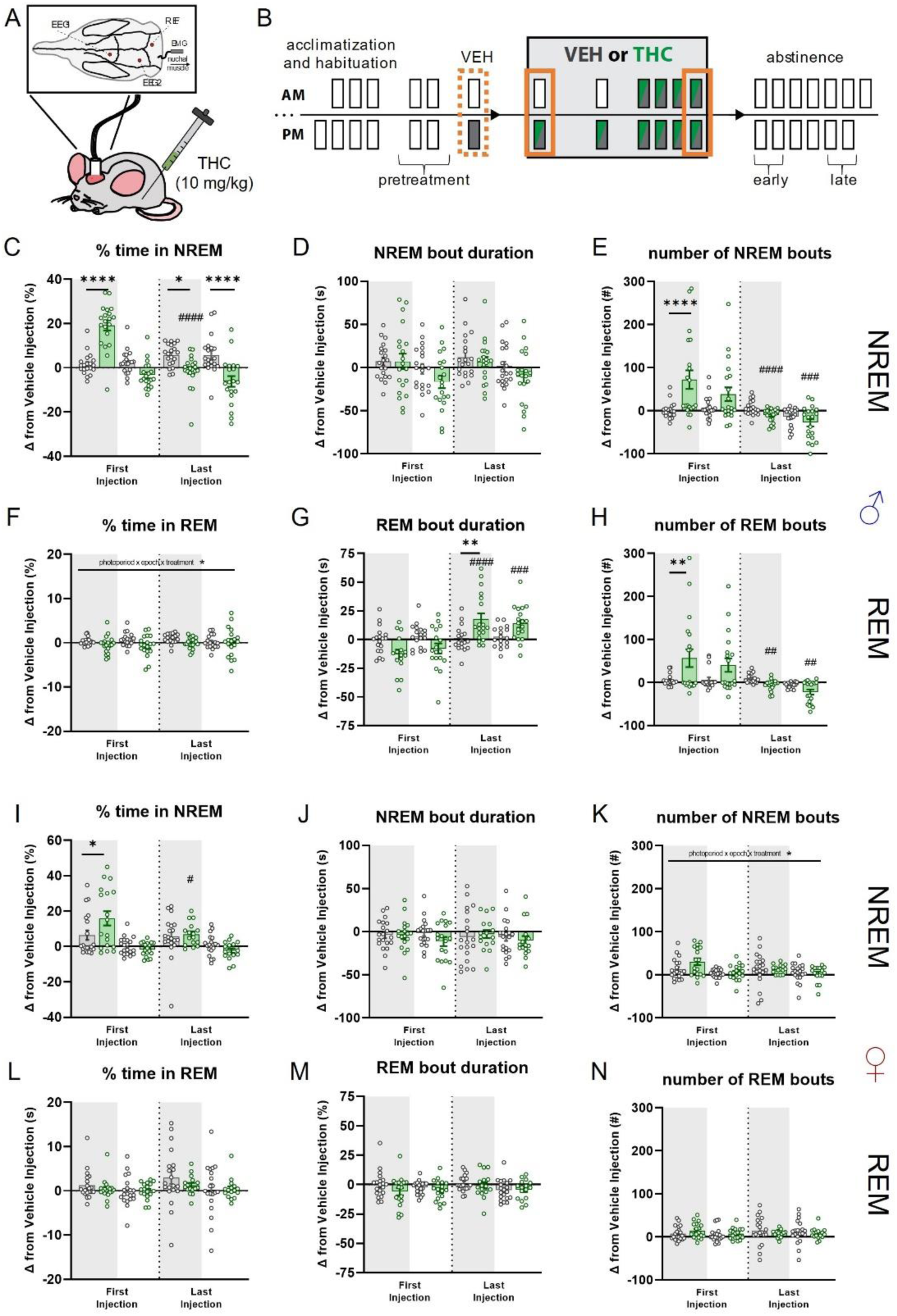
Effects of acute and chronic THC administration on sleep in male and female mice. (A) Cartoon illustrating i.p. administration of 10mg/kg THC in chronically tethered mice implanted with two EEG, one EMG, and one reference electrodes and their relative placements on the skull / nuchal tissue. (B) Timeline illustrating treatment regimen and epochs contributing to analysis. Dotted orange line indicates vehicle treatment epoch that acute and chronic administration epochs are compared. (C) Comparison between effects of THC or VEH on changes from vehicle injection in percent time spent in NREM sleep in male mice. There is a significant interaction (F_epoch x treatment_(1,38) = 66.18; p = <0.0001), with post-hoc tests finding differences between-groups during First Injection-LOFF (p**** = <0.0001), Last Injection-LOFF (p* = 0.0346) and LON (p**** = <0.0001). The THC group had significant within-group difference between First and Last Injection LOFF (p^####^ = <0.0001). (D) Comparison between effects of THC or VEH on changes from vehicle injection in NREM bout duration in male mice. No significant effects were detected. (E) Comparison between effects of THC or VEH on changes from vehicle injection in number of NREM bouts in male mice. There is a significant interaction (F_epoch x treatment_(1,38) = 11.68; p = 0.0015), with post-hoc tests finding differences between-groups during First Injection-LOFF (p**** = <0.0001). The THC group had significant within-group difference between First and Last Injection LOFF (p^####^ = <0.0001) and LON (p^###^ = 0.0001). (F) Comparison between effects of THC or VEH on changes from vehicle injection in percent time spent in REM sleep in male mice. There is a significant interaction (F_photoperiod x epoch x treatment_(1,30) = 4.573; p = 0.0407). (G) Comparison between effects of THC or VEH on changes from vehicle injection in REM bout duration in male mice. There is a significant interaction (F_epoch x treatment_(1,36) = 27.37; p = <0.0001), with post-hoc tests finding differences between-groups during Last Injection-LOFF (p** = 0.0112). The THC group had significant within-group difference between First and Last Injection of LOFF (p^####^ = <0.0001) and LON (p^###^ = 0.0002). (H) Comparison between effects of THC or VEH on changes from vehicle injection in number of REM bouts in male mice. There is a significant interaction (F_epoch x treatment_(1,36) = 9.043; p = 0.0048), with post-hoc tests finding differences between-groups during First Injection-LOFF (p** = 0.0051). The THC group had significant within-group difference between First and Last Injection LOFF (p^##^ = 0.0011) and LON (p^##^ = 0.0013). (I) Comparison between effects of THC or VEH on changes from vehicle injection in percent time spent in NREM sleep in female mice. There is a significant interaction (F_photoperiod x treatment_(1,38) = 5.881; p = <0.0202), with post-hoc tests finding differences between-groups during First Injection-LOFF (p8 = 0.0215). The THC group had significant within-group difference between First and Last Injection LOFF (p^#^ = 0.0422). (J) Comparison between effects of THC or VEH on changes from vehicle injection in NREM bout duration in female mice. No significant effects were detected. (K) Comparison between effects of THC or VEH on changes from vehicle injection in number of NREM bouts in female mice. There is a significant interaction (F_photoperiod x epoch x treatment_ (1,25) = 6.629; p = <0.0163). (L) - (N) Comparison between effects of THC or VEH on changes from vehicle injection in percent time in REM (L), bout duration (M), and number of bouts (N) in female mice. No significant effects were detected.

Small changes in percent time in REM sleep were observed following the first THC injection, but post-hoc testing found no significant differences between treatments or within treatment groups across epochs (**Figure 2F**). In contrast, REM bout duration, particularly after the last injection where THC treated male mice showed larger increases compared to VEH treated mice during LOFF. THC treated male mice also had longer REM bout durations after the last injection compared to first injection during both photoperiods (**Figure 2G**). THC treated male mice also exhibited increased number of REM bouts during LOFF after the first injection compared to VEH treated mice, while the number of REM bouts in these THC treated decreased after the last injection when compared to the first injection for both LOFF and LON (**Figure 2H**).

Female mice also exhibited increased percent NREM sleep after acute THC (**Figure 2I**), that was abolished after repeated dosing. In contrast to male mice (**Figure 2C**), NREM sleep time did not differ between THC treated females and the VEH control group after repeated dosing. Neither acute nor chronic THC administration altered NREM bout duration in female mice (**Figure 2J**). While there was an overall increase in the number of NREM bouts, there were no pair-wise differences between groups (**Figure 2K**). Also, in contrast to male mice, we observed no effect of THC injections on REM sleep time or architecture (**Figure 2L-N**).

#### Chronic THC transiently disrupts sleep in male mice at early abstinence timepoints

Because sleep disturbances are commonly reported by individuals with CUD during periods of abstinence [35, 57, 58], we obtained electrographic measures of sleep after the six-day chronic treatment regimen and normalized these to the pretreatment baseline for each subject (**Figure 3A and B**). In male mice, we observed reduced NREM sleep following chronic THC treatment, evidence of acute, spontaneous withdrawal symptoms in early abstinence (**Figure 3C**). These effects were specific to LOFF in early abstinence, and they recovered by late abstinence, when THC-treated males slept no differently than the VEH group. Reduced sleep in early abstinence was largely due to a reduction in the duration of NREM bouts with THC-treated subjects exhibiting decreased bout duration during LOFF in early but not late abstinence (**Figure 3D**). While interaction effects for NREM bout number were observed in late-LOFF compared to early-LOFF in male THC treated mice, post-hoc tests found no significant differences related to treatment (**Figure 3E**).

**Figure 3.**
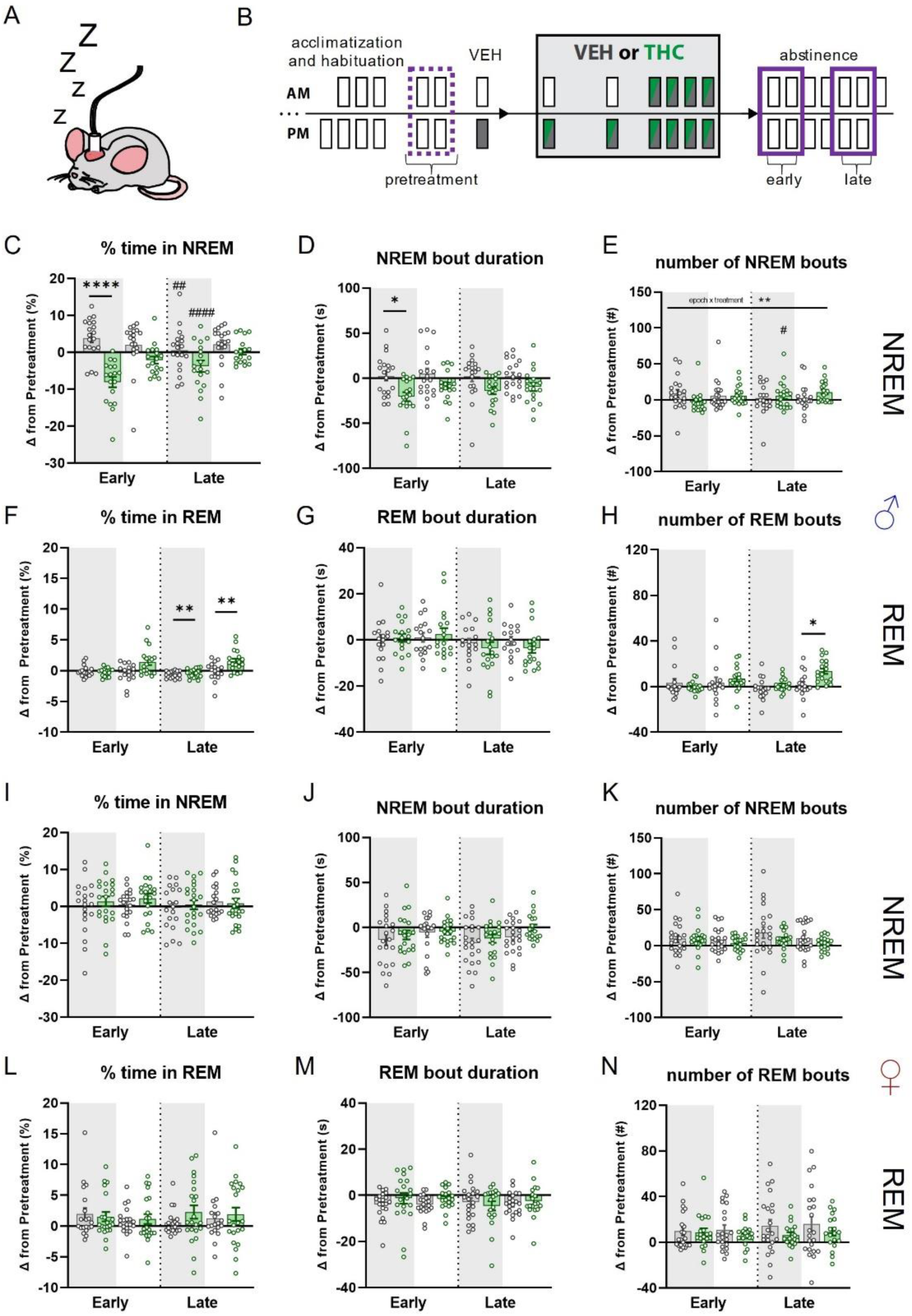
Effects of chronic THC administration on sleep in male and female mice during early and late abstinence. (A) Cartoon illustrating chronically tethered mouse during epochs where no injection is given, i.e., pretreatment and abstinence. (B) Timeline illustrating treatment regimen and epochs contributing to analysis. Dotted purple line indicates average of two pretreatment epochs that average of days 1 and 2 of early and days 5 and 6 of late abstinence are compared. (C) Comparison between effects of THC or VEH on changes from pretreatment in percent time spent in NREM sleep in male mice. There is a significant treatment effect (F_treatment_(1,35) = 20.64; p = <0.0001), with post-hoc tests finding differences between-groups during Early-LOFF (p**** = <0.0001). Both groups had significant within-group difference between Early and Late abstinence LOFF (THC p^####^ = <0.0001; VEH p^##^ = 0.0030). (D) Comparison between effects of THC or VEH on changes from pretreatment in NREM bout duration in male mice. There is a significant treatment effect (F_treatment_(1,35) = 11.14; p = 0.0020), with post-hoc tests finding differences between-groups during Early-LOFF (p* = 0.0113). (E) Comparison between effects of THC or VEH on changes from pretreatment in number of NREM bouts in male mice. There is a significant interaction (F_epoch x treatment_(1,37) = 12.64; p = 0.0011). (F) Comparison between effects of THC or VEH on changes from pretreatment in percent time spent in REM sleep in male mice. There is a significant treatment effect (F_treatment_(1,33) = 8.048; p = 0.0077), with post-hoc tests finding differences between-groups during Late-LOFF (p** = 0.0017) and Late-LON (p** = 0.0057). (G) Comparison between effects of THC or VEH on changes from vehicle injection in REM bout duration in male mice. No significant effects were detected. (H) Comparison between effects of THC or VEH on changes from pretreatment in number of NREM bouts in male mice. There is a significant interaction (F_epoch x treatment_(1,33) = 5.744; p = 0.0224), with post-hoc tests finding differences between-groups during Late-LOFF (p* = 0.0228). (I) - (N) Same metrics as (C)-(H) but in female mice. No significant effects were detected.

Chronic THC treatment in male mice led to increased time spent in REM sleep compared to VEH treatment in LOFF and LON periods late in abstinence (**Figure 3F**). We did not detect any treatment related effects in REM bout duration (**Figure 3G**). THC treated male mice had greater increases in REM bout number when compared to VEH treated mice during LOFF in late abstinence (**Figure 3H**). In contrast to male mice, no significant treatment-related main or interaction effects were detected for NREM or REM metrics during abstinence in females (**Figure 3I-N**).

#### Low dose chronic treatment does not strongly alter sleep during administration or abstinence

We next performed separate experiments where mice were chronically treated with a lower dose (1.0 mg/kg) of THC. (**Supplemental Figure 2A & B**). During the treatment period we only detected a decrease in NREM bout duration and number in THC treated relative to VEH treated mice during LON after the last injection (**Supplemental Figure 2C-E**). A significant interaction in number of REM bouts was detected, but no significant differences between treatments or across injection epochs was found in post-hoc tests (**Supplemental Figure 2F-H**). We found no significant main or interaction effects of low dose chronic THC treatment in female mice for NREM sleep metrics, nor percent time in REM or number of REM bouts. We did detect an interaction effect on REM bout duration in these groups, but post-hoc tests found no significant treatment-related effects (**Supplemental Figure 2I-K**). Low dose chronic THC treatment did not alter any sleep metrics during early or late abstinence when compared to the pretreatment epoch in either sex (**Supplemental Figure 3A-N**).

#### Acute THC treatment does not alter sleep during abstinence

Profound sleep disturbances were only observed in male mice after chronic 10 mg/kg THC treatment. These changes could be due to effects of the final THC treatment in the chronic regimen rather than changes accrued during chronic treatment. To examine this possibility, we gave a single 10 mg/kg THC treatment and examined sleep in abstinence. A significant interaction was detected for changes in NREM and REM bout duration, but no significant differences between treatments or across abstinence epochs were observed (**Supplemental Figure 4A-H**). Thus, the effects of chronic 10 mg/kg THC in male mice are likely due to accrued effects of chronic treatment.

### Sustenance intake and locomotor activity during chronic THC treatment and abstinence

Accumulating evidence indicates humans experience altered hunger and thirst during cannabis withdrawal. These symptoms are often accompanied by restlessness which might manifest as increased locomotor behavior in general contexts. Thus, we examined how food and water intake and locomotor behavior are altered during early and late abstinence following chronic THC treatment in mice (**Figure 4A-B**). For both males and females, significant treatment related effects were detected for food and water intake, and locomotion (**Figure 4C-H**). Additionally, THC treated animals showed decreased food and water intake during LOFF after the first injection of the chronic treatment regimen, which differed from VEH treated mice. Changes from vehicle treatment did not differ between groups after the last chronic treatment, but after the last injection both sexes showed normalization of intake back towards vehicle treatment levels during LOFF. Similarly, THC treated mice of both sexes had reduced changes in locomotion compared to VEH treated mice during LOFF after the first treatment. While VEH treated mice showed reduced locomotion after the last treatment compared to first treatment, an increase was found in THC treated mice. THC treatment did not alter food or water intake and locomotion during early or late abstinence when compared to the pretreatment epoch in either sex (**Figure 4I-N**).

**Figure 4.**
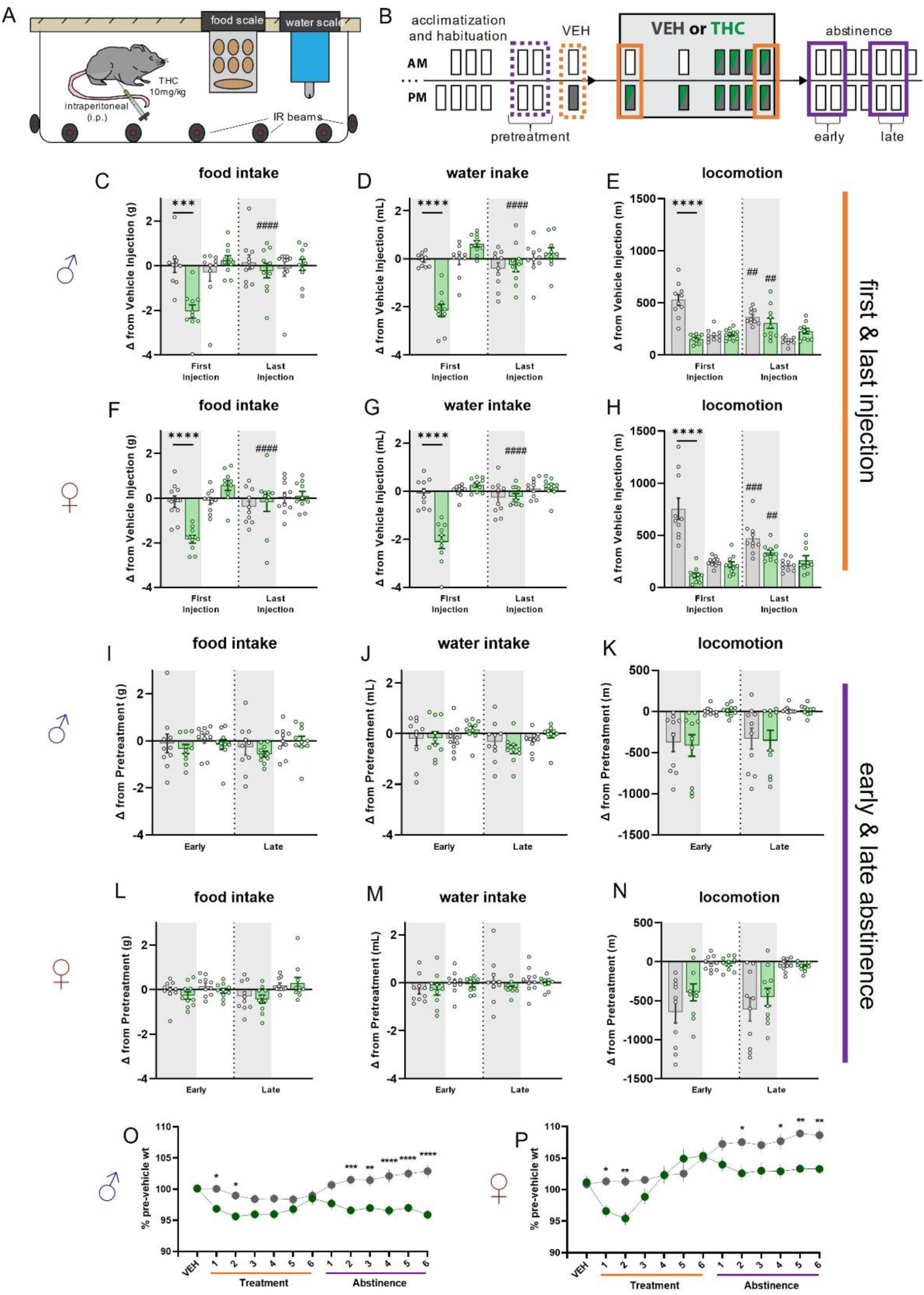
Measurements of changes in food and water intake, and locomotion, during first and last THC injections and early and late abstinence. (A) Cartoon illustrating home-cage intake and locomotion monitoring platform. (B) Timeline illustrating treatment regimen and epochs contributing to analysis. Dotted orange line indicates vehicle treatment epoch that acute and chronic administration epochs are compared, and dotted purple line indicates average of two pretreatment epochs that average of days 1 and 2 of early and days 5 and 6 of late abstinence are compared. (C) Comparison between effects of THC or VEH on changes from vehicle injection in food intake in male mice. There is a significant interaction effect (F_epoch x treatment_(1,18) = 11.37; p = 0.0034), with post-hoc tests finding differences between-groups during First Injection-LOFF (p*** = 0.0007) and within group differences for THC treated mice during Last Injection LOFF (p^####^ = <0.0001). (D) Comparison between effects of THC or VEH on changes from vehicle injection in water intake in male mice. There is a significant interaction effect (F_epoch x treatment_(1,18) = 14.43; p = 0.0013), with post-hoc tests finding differences between-groups during First Injection-LOFF (p**** = <0.0001) and within group differences for THC treated mice during Last Injection LOFF (p^####^ = <0.0001). (E) Comparison between effects of THC or VEH on changes from vehicle injection in locomotion in male mice. There is a significant treatment effect (F_treatment_(1,18) = 11.42; p = 0.0033), with post-hoc tests finding differences between-groups during First Injection-LOFF (p**** = <0.0001) and within group differences for THC treated (p^##^ = 0.0037) and VEH treated (p^##^ = 0.0013) mice during Last Injection LOFF. (F) Comparison between effects of THC or VEH on changes from vehicle injection in food intake in female mice. There is a significant interaction effect (F_epoch x treatment_(1,18) = 4.964; p = 0.0389), with post-hoc tests finding differences between-groups during First Injection-LOFF (p**** = <0.0001) and within group differences for THC treated mice during Last Injection LOFF (p^####^ = <0.0001). (G) Comparison between effects of THC or VEH on changes from vehicle injection in water intake in female mice. There is a significant treatment effect (F_treatment_(1,18) = 18.517; p = 0.0092), with post-hoc tests finding differences between-groups during First Injection-LOFF (p**** = <0.0001) and within group differences for THC treated mice during Last Injection LOFF (p^####^ = <0.0001). (H) Comparison between effects of THC or VEH on changes from vehicle injection in locomotion in female mice. There is a significant treatment effect (F_treatment_(1,18) = 21.05; p = 0.0002), with post-hoc tests finding differences between-groups during First Injection-LOFF (p**** = <0.0001) and within group differences for THC treated (p^##^ = 0.0026) and VEH treated (p^###^ = 0.0002) mice during Last Injection LOFF. (I) - (N) Same metrics as above but for early and late abstinence epochs. No significant effects were detected for either sex. (O) Percent of Vehicle treatment weights during chronic THC treatment and abstinence in male mice. There is a significant treatment effect (F_treatment_(1,18) = 11.08; p = 0.0037), with post-hoc tests finding differences between-groups (p* = < 0.05, p** = < 0.01, p*** = <0.001, p**** = <0.0001). (P) Percent of Vehicle treatment weights during chronic THC treatment and abstinence in female mice. There is a significant treatment effect (F_treatment_(1,18) = 5.577; p = 0.0297), with post-hoc tests finding differences between-groups (p* = < 0.05, p** = < 0.01).

THC treated mice of both sexes showed significantly lower body weight compared to VEH treated mice during the first two days of treatment. This effect normalized by the final THC treatment but reemerged during abstinence (**Figure 4O-P**).

### Altered reward seeking and conditioned cue discrimination during chronic THC abstinence

We hypothesized that various aspects of reward seeking and consumption would be altered during THC abstinence (**Figure 5A**), consistent with observed alterations in striatal DA release and sleep disturbances. We examined these behaviors by training mice to respond on levers that led to the presentation of auditory stimuli signaling whether-or-not sucrose-solution reward would be delivered (**Figure 5B-C**). Over 5 pretraining sessions, all mice displayed increased latency to enter the sucrose port upon lever press-contingent presentation of an auditory stimulus signaling no reward (CS-), while maintaining relatively short latency when presented with the stimulus signaling the reward (CS+) (**Figure 5D**) indicating that the mice learned to discriminate CS+ from CS-.

**Figure 5.**
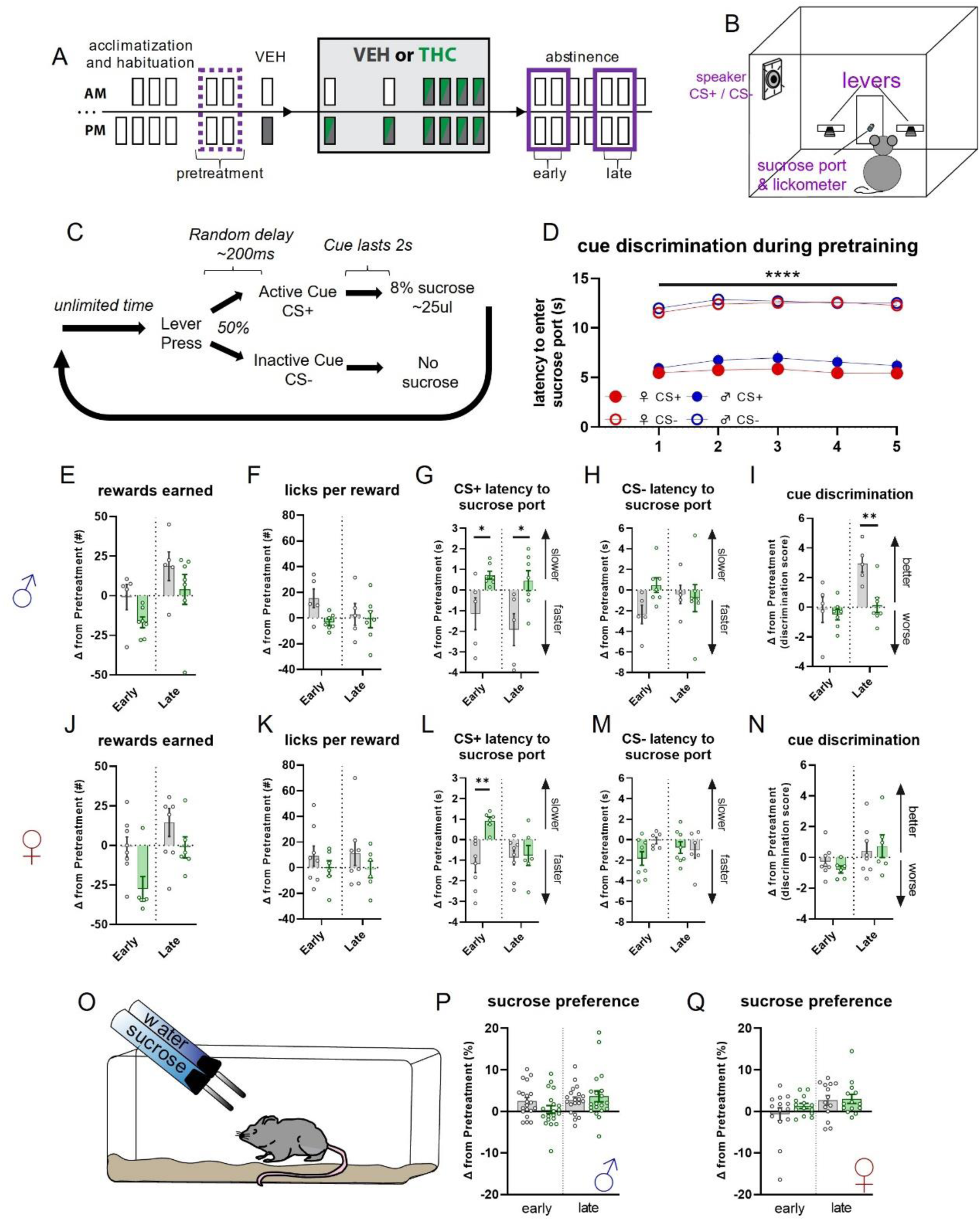
Changes in reward seeking behaviors during early and late abstinence from chronic THC treatment in male and female mice. (A) Timeline illustrating treatment regimen and epochs contributing to analysis. Dotted purple line indicates average of two pretreatment epochs that average of days 1 and 2 of early and days 5 and 6 of late abstinence are compared. (B) Cartoon illustrating operant chamber for reward seeking and conditioned auditory cue presentation. (C) Diagram showing operant sucrose-seeking procedure with cue discrimination. (D) Cue discrimination during last five pretraining sessions as measured by latency to enter reward port. There is a significant effect of cue (2_cue_ ×5_session_ RM-ANOVA; F_cue_(1,45) = 182.9; p = <0.0001). (E) - (I) Changes in behavioral metrics related to reward seeking and cue discrimination in male mice. There is a significant treatment effect for change in latency to approach sucrose port after CS+ (G; F_treatment_(1,10) = 8.616; p = 0.0149) with post-hoc tests finding group differences during Early (p* = 0.0261) and Late (p* = 0.0131) abstinence, and significant treatment effects in cue discrimination (I; F_treatment_(1,10) = 5.830; p = 0.0364) with posthoc tests finding group differences during Late abstinence (p** = 0.0045). (J) - (N) Same as E-I but for female mice. There is a significant treatment effect for change in latency to approach sucrose port after CS+ (L; F_treatment_(1,12) = 5.787; p = 0.0332) with post-hoc tests finding group differences during Early abstinence (p** = 0.0014). (O) Cartoon depicting sucrose preference test. (P) - (Q) Change in sucrose preference score during Early and Late abstinence in male (P) and female (Q) mice. No significant effects were detected.

Comparing behavior during pretreatment and following chronic THC or VEH treatment indicated no significant differences in rewards earned (**Figure 5E**). THC treatment did not alter sucrose consumption behavior (i.e., ‘licks’) upon reward delivery (**Figure 5F**). The most striking effect was the behavioral response to reward predictive cues, where THC treated mice were slower to approach the sucrose reward port after a lever press indicating reward availability, i.e., CS+ latency (**Figure 6G**), in both early and late abstinence. This effect was not present for non-rewarded (i.e., CS-) lever press latency (**Figure 5H**). Interestingly, THC treated male mice also differed from VEH treated mice in their change in cue discrimination during late withdrawal (**Figure 5I**). Here, the VEH mice increased the ratio of CS-latency to CS+ latency, while the THC mice showed ratios similar to pretreatment and early withdrawal levels, indicating THC treatment attenuated the enhanced discrimination between the cues that VEH treated mice experience during late withdrawal. Like male mice, THC treated female showed no significant differences in rewards earned (**Figure 5J**), or reward consummatory behaviors (**Figure 5K**), and had slower CS+ latency during early withdrawal (**Figure 5L**) with no difference in CS-latency (**Figure 5M**). Unlike males, no effect on cue discrimination was observed in female mice for either abstinence epoch (**Figure 5N**).

To examine whether differences in reward seeking behaviors were due to altered hedonic characteristics of sucrose reward, we tested mice in a sucrose preference test (**Figure 5O**). No differences in sucrose preference during early or late abstinence were observed for THC treated mice of either sex compared to VEH treated mice (**Figure 5P-Q**).

### Irritability measurements and circulating stress hormones

Clinical evidence suggests some cannabis users display enhanced aggression and irritability during abstinence [13, 20, 59]. We used the bottle brush test [60, 61] (**Supplemental Figure 5A**) to assess changes in irritability-related behaviors during abstinence in chronic THC vs VEH treated mice (**Supplemental Figure 5B**).

#### Sex differences emerge during ethological dissection of irritability behaviors

We scored metrics related to aggressive (smelling, biting, boxing, following, exploring, tail rattling) and defensive/stress-induced behaviors (escaping, digging, jumping, climbing, defecating, vocalizing, grooming) (**Supplemental Figure 5C-P**). Male THC treated mice showed enhanced boxing behavior during both early and late abstinence (**Supplemental Figure 5D**). In addition, male THC but not VEH treated mice had a greater change in escape behaviors in late as compared to early abstinence (**Supplemental Figure 5F**). No significant changes in irritability behaviors were detected in female mice. Biting, exploring, tail rattling, jumping, climbing, and vocalizing were scored, but were scarce or absent in most mice, so were not further analyzed.

#### THC withdrawal symptoms are not likely driven by changes in stress as measured by circulating corticosterone

Stress is known to have profound effects on sleep and behavior [62]. We assessed whether the THC treatment itself acted as a stressor in a manner that might alter sleep and wake behaviors during abstinence. We measured circulating plasma corticosterone (CORT) as an index of perceived stress and assessed changes in pretreatment CORT during early and late withdrawal (**Supplemental Figure 6A**). While THC treated male mice displayed increased CORT during early withdrawal compared to VEH treated mice (**Supplemental Figure 6B**), this increase (27.26 ng ± 13.17 ng; mean ± SEM) was an order of magnitude smaller than CORT increases from chronic mild stress [63]. Changes in CORT did not differ between THC treated and VEH treated female mice (**Supplemental Figure 6C**).

## Discussion

### Summary

We examined how spontaneous withdrawal from chronic THC administered to naïve mice drives changes in striatal DA release, sleep disruptions, and behavioral maladaptation. Overall, our experiments show that mice can be used to study translatable THC withdrawal symptoms and neural mechanisms that might drive these symptoms. Of particular interest are apparent sex-differences in nearly all metrics during spontaneous THC withdrawal. These studies open the door for further investigation into the neurophysiological changes mediating CWS and potential therapeutic interventions to treat CUD/CWS. Key findings and implications are discussed below.

Striatal DA has well established roles in drug dependence [52], including CUD [51, 64], and DA dysregulation following chronic cannabis or THC use is a known factor in the manifestation of CWS [65]. The implications of the sex differences in DA release during abstinence after acute THC are unclear. Perhaps the increased DA release in males makes them susceptible to enhanced sensitivity to salient or rewarding stimuli after their first drug experience, while females are resistant to such changes. Future studies are required to test whether the increased DA release in males contributes to behavioral and sleep alterations following chronic THC administration.

More pertinent to CUD/CWS are the observed sex differences in DA release during abstinence after chronic THC treatment. In early abstinence female mice had far greater DA release across the striatum, while males showed no such effects. However, while in females these measurements returned to control level in late abstinence, the relative lack of change in DA release in males may result in susceptibility to changes in reward-associated stimuli. These findings may be related to effects of altered dopaminergic transmission in disrupted cognitive function due to dysregulation in signal to noise processing [66]. Again, further investigation into the role of altered DA release in wake behaviors and sleep is needed.

Previous studies have mainly examined effects of chronic cannabis or THC on firing of ventral tegmental area DA neurons that project to striatum, with less information about changes during abstinence. Furthermore, it is unclear how these firing changes relate to alterations in striatal DA release, as that has not been measured in past studies. Diana and coworkers found that rat midbrain DA single-unit activity was reduced 24hrs after the last THC injection using a treatment regimen similar to the one we used [67]. Based on this finding one might not expect the increased striatal DA release that we observed during abstinence. A possible explanation for this contradiction is the recent evidence that striatal DA activity does not necessarily reflect midbrain DA neuron activity [68], and this dissociation can be explained by intrastriatal microcircuitry. Indeed, the endocannabinoid (eCB) system can modulate striatal DA release independent of postsynaptic action potentials [69], suggesting prolonged alterations in striatal eCB action due to THC administration as a potential contributor to the increased DA release. Imaging studies in human cannabis users abstaining for 1-7 days found a reduction in stimulant-induced DA response [65], which also contrasts with our findings of increased DA release at similar timepoints during withdrawal. Technical differences can potentially reconcile these divergent findings, as stimulant-mediated increases involve altered DA transporter function while electrical stimulation induces DA release directly via DA afferent stimulation and indirectly via cholinergic interneuron activation [70]. Examining the role of cholinergic drive on DA release may thus also indicate mechanisms underlying THC actions. Likewise, our data were collected postmortem in brain slice preparations taken at precise withdrawal timepoints. Future in-vivo studies using genetically coded DA sensors [71] can also shed light on the relationship between striatal DA activity and precise THC withdrawal timepoints.

### A mouse model for sleep disruption during THC withdrawal

Treating sleep pathologies is often given as a reason for THC use both recreationally and medically [72, 73]. Indeed, many individuals list poor sleep as a major factor leading to their relapse to cannabis use [16], and those subjects showing sleep disruption, poor sleep quality in particular, relapse more readily [26, 27, 74, 75]. However, sleep disruption is one of the most consistent and problematic aspects of cannabis withdrawal, with altered sleep observed in the majority of regular users who attempt to quit [16, 21].

The observed effects on sleep, especially male mice are generally consistent with our hypothesis that chronic THC administration would cause sleep disruption that diminishes as mice build tolerance to the drug. However sleep disruption would reemerge early in withdrawal and normalize during abstinence – i.e. the disruption would mirror the transient changes observed in the clinical setting [17]. The observations that acute THC treatment enhanced NREM and REM sleep was fragmented in male mice during abstinence following acute THC are consistent with previous reports of single THC or cannabis exposures in animal models [76–78] and humans [32, 79–83]. The finding that THC tolerance following chronic treatment diminishes the drug’s effect on sleep is also consistent with past findings in animals [77, 78, 84–86] and regular human cannabis users [34, 79, 82, 87–89]. We also found that male mice experience disruptions during THC withdrawal similar to human cannabis users [35, 36, 57, 90, 91] while female mice appear more resilient to the overall effects of spontaneous THC withdrawal on sleep. The absence of significant differences in female mice in our study was surprising given that female cannabis users report sleep difficulty during cannabis abstinence [92]. However, in clinical polysomnographic studies measuring sleep during cannabis abstinence the participants are primarily male, making comparison to our findings in female mice difficult. Future clinical studies should be properly powered to examine sleep physiology in female cannabis abstainers. Additionally, if female cannabis users are indeed more resilient to sleep changes as measured by polysomnography, there are other psychological manifestations occurring during cannabis or THC withdrawal that may drive self-report of poor sleep quality despite no profound changes in sleep physiology.. We believe overall our study provides a valuable back-translational model to investigate the neural mechanisms mediating sleep disruption during cannabis withdrawal.

### Mouse as a back-translational model for CWS during wake-behaviors

Past research using mice to study spontaneous CWS primarily report negative findings for behaviors beyond typical somatic symptoms and cannabis tetrad metrics [3]. This is potentially confounded by typical behavioral paradigms used in preclinical settings that may not adequately probe the most pronounced human CWS. We focused our behavioral tasks to measure changes in behaviors that directly map to some of the strongest CWS; (1) reduced food and water intake, (2) restlessness (i.e., heightened locomotion), (3) amotivation, (4) attention deficits, and (5) irritability.

While we observed profound hypophagia, hypodipsia, and reduced locomotion behavior after the first THC injection, our chronic THC treatment clearly induces behavioral tolerance to the effects of the drug as evidenced by the normalization of these measures to control levels by the final THC injection. However, the lack of change in these metrics in abstinence following chronic THC exposure suggest home cage intake and locomotion may not serve as an appropriate back-translational model for human CWS. Perhaps this paradigm does not recapitulate other environmental and cognitive factors that manifest in the human population, e.g., additional life stressors, or situations that when compounded with cannabis abstinence affect food and water intake. Nonetheless, we observed significant differences between treatment groups in body weight as early as day two of abstinence. This finding is particularly intriguing as it suggests chronic THC treatment alters metabolic function that may contribute to alterations in body weight despite no differences in intake or locomotion – a notion that is ripe for future investigation.

Our observation that male and female mice did not differ in rewards earned indicates that CWS in mice does not involve strong changes in motivation or drive. However, the slower retrieval of reward in THC treated mice of both sexes may indicate decreased motivation. Responses to the CS+ and CS-cues that followed operant responding provide insight into attention during this goal-directed task. In particular, the poorer cue discrimination in male mice treated with THC in late abstinence indicates a deficit in attention or focus that emerges later in withdrawal. While this operant responding paradigm provided evidence of impairment in several cognitive processes related to goal-directed action during chronic THC withdrawal, future studies should directly target specific aspects of reward-seeking behaviors and attention processes in an effort to home in on neural mechanisms mediating these specific disruptions.

Increased irritability is prevalent during cannabis abstinence in humans [24, 93]. We were surprised to find that relatively few irritability-like behaviors were altered in our bottle brush test assay. Nonetheless, male mice showed enhancement of some irritability-like metrics, i.e., boxing and escape behaviors. These behaviors are interesting in that they demark disparate types of irritability behaviors – aggressive and defensive, respectively, indicating that male mice experience alterations in irritability behavior in general, though these alterations manifest in somewhat specific behavioral outputs in the bottle brush test.

### Conclusions and Caveats

Technical limitations to our studies, primarily that metrics were gathered from separate cohorts of animals, make it difficult to draw conclusions about correlations between various behaviors and, in particular, DA voltammetry measurements. Nonetheless, a general observation is that female mice appear more resilient to effects of chronic THC treatment on sleep and maladaptive behaviors during abstinence. A speculative explanation is the elevation in striatal DA release we observe in female mice during early withdrawal may play a protective role in attenuating withdrawal symptoms, and the clearest indicator of this effect in our studies is in the relationship between lack of sleep disruption and striatal DA release in early abstinence in female mice. Striatal DA is known to play a role in sleep-state architecture [46], so further studies are needed to investigate the causal role of enhanced striatal DA release observed in female mice in alleviating sleep disruption during early chronic THC abstinence.

Our findings provide a model of significant and translatable CWS in rodents using a protocol that does not require antagonist-precipitated withdrawal. Future studies based on the treatment protocol and time points used in our study can unravel the likely complex neural mechanisms that drive sleep disruption and behavioral maladaptation that are prevalent in CWS – crucial first steps towards therapeutics to combat CUD.

## Funding

This work was supported by the Division of Intramural Clinical and Biological Research of the National Institute on Alcohol Abuse and Alcoholism [grant number ZIA000416 to DML].

## Author contributions

AJK, KPA, MJP, YM, and DML designed experiments. AJK, KPA, MJP, SRM, ALG, and HBC performed behavioral experiments. YM and RTP performed FSCV experiments. AJK, KPA, and MJP performed statistical analysis of data. AJK wrote initial draft of manuscript, and after which all authors discussed the results and contributed to the final manuscript.

## Declaration of competing interest

The authors declare no competing interests or conflicts of financial interest.

## Supplemental Methods and Materials

### Subjects

C57Bl6/J mice were obtained from the Jackson Laboratory (Bar Harbor, ME), and housed 2 to 4 per cage. Initially, mice weighed 25 to 30 g, and their body weight did not change substantially throughout the study. Upon arrival to the animal colony, all mice were maintained on a 12:12 h light:dark cycle with lights turning on at 06:30 EST and off at 18:30 EST. Throughout all parts of the study, except during hour long operant responding experiments, subjects were provided with ad libitum access to food and water and housed in an environment maintained at 22.2 °C and 50% humidity. The experimenter conducting experiments where data was collected was blind to experimental treatment groups, except for measurement of weights during chronic treatment administration.

### Sleep recordings

#### Surgical preparation for sleep recordings

Surgeries were performed on subjects anesthetized with isoflurane. Custom electrode implants were prepared as previously described [1]. Stereotaxic surgery was performed to implant electrodes over medial frontal cortex (AP: +1.50, ML: ±1.00) and parietal cortex (AP: −2.50, ML: ±2.00). A ground/reference electrode was also implanted over the cerebellum (AP: −5.7, ML: ±1.7). Electrodes consisted of stainless-steel screws (AMS90/1P-25; Antrin Miniature Specialties Inc., Fallbrook, CA) placed in contact with the brain surface and wrapped with Teflon-insulated fine stainless-steel wires (Cat # 91400; A-M Systems Inc., Sequm, WA), that in turn connected to a headcap (MS363 & E363-0; Plastics One, Roanoke, VA) for later connection with recording tethers (363-363; Plastics One, Roanoke, VA). A 1.25mm stainless steel suture pad also connected to the headcap was inserted beneath the nuchal muscle before the entire headcap was fixed to the skull using standard cold-cure dental acrylic. Post-operative pain and discomfort were mitigated by administering ketoprofen (5 mg/kg i.p.) analgesic solution immediately after the surgery was completed and once daily for the next two days. Following surgery, subjects were allowed to recover for two weeks before being tethered to recording equipment.

#### Recording environment

After the recovery period mice were brought to an isolated room dedicated to 24-hour, long term, mouse sleep recording experiments. Subjects were placed individually into recording cages (custom fabricated, 4 L polycarbonate buckets, Cambro RFSCW4135, Webstaurant Store, Lancaster, PA) and tethered to commutators (SL6C/ SB; Plastics One, Roanoke, VA). These cages were placed into electromagnetically shielded chambers that provided sound and light attenuation (SAE chambers; 5 cages/chamber; CT-ENV-018MD-EMS-X1, MED Associates, Fairfax, VT), with five individual cages present in each chamber such that mice could see, smell, and hear one another. The chambers were outfitted with small fans which continuously circulated room air and provided stable background noise within the recording chambers, and white LED light strips (#10434, General Electric, Fairfield, CT) controlled by a timer synchronized with ambient room lighting. Standard food pellets (BioServe; Cat no. NIH31) were placed on the cage bedding and access to water was provided via glass tubes (#9019, Bio-serve, Frenchtown, NJ). White noise generators were continuously running in the recording room itself to further mask outside noise from disturbing mice.

#### Polysomnographic (EEG/EMG) recording

Recordings of polysomnographic signals (EEG/EMG) were obtained from the chronically tethered mice over 23hr periods. During the intervening 1hr, data acquisition was halted to check on the animals to ensure ad libitum food and water were available and perform injections of THC or vehicle when appropriate. EEG/EMG signals were amplified 1000x (20x HST/16V-G20 head-stage followed by 50x wide-band PBX, Plexon, Dallas, TX) and digitized at 1 kHz (PCI-6071E, National Instruments, Austin, Tx). Data collection used a standard PC computer (Optiplex GX620, Dell Computers, Round Rock, Tx) running Recorder v2 software (Plexon). A 60 Hz digital notch filter was applied to attenuate line noise on all channels. EEG channels were then low-pass filtered at 120 Hz with a 2-pole Bessel filter, and EMG channels were high-pass filtered 40 Hz with a 4 pole Bessel filter. Data were saved to disk for analysis offline.

#### Scoring of vigilance states and sleep architecture

##### Constructing the state space

EEG/EMG signals were analyzed offline to categorize epochs of wake, NREM, and REM states using software written int MATLAB and CUDA C. This methodology has been previously validated in our laboratory [1] and has been used in several sleep-wake architecture analysis [1, 2]. The following analysis pipeline was conducted for each mouse for each recording session. Briefly, data were scored in 2 s epochs by clustering power spectral features in a 3-dimensional state space. The three power spectral features that made up the state space orthogonal vectors were: (1) a power spectral ratio (0.5–20 Hz/0.5–100 Hz) from the frontal EEG channel, known to separate NREM from wake and REM [3]; (2) a second power spectral ratio from the occipital EEG channel to compute the prominence of theta and low alpha relative to delta power (5–10 Hz/0.5–4 Hz). This ratio has been used to help separate REM epochs which are characterized by an overall increase in the theta bandwidth [4–6]; and (3) the root mean square values of the EMG power spectra. Finally, each of these state space vectors was smoothed by convolution via a 10 s Hann window, median-centered, and normalized to the maximum absolute value.

##### Vigilance state classification

A three-step process was implemented to categorize vigilance state once the state space was calculated [1]. First, a rough estimate of the boundaries for each state based on univariate distributions along each axis were used to produce a starting point for establishing a preliminary score. Next, a 3-dimensional product kernel estimate using a Gaussian kernel function was calculated for each cluster (i.e., wake, NREM, REM, and unassigned) and these estimates were scaled such that their maximal values were equal to the value in the corresponding peak of the overall density function. We established 99% confidence intervals from this estimate that formed inclusion criteria for the three clusters. Finally, we applied a transitional classifier to clean up unassigned points that were bounded by epochs of the same state. The scoring results for each subject were visually inspected to ensure accuracy, and subjects with excessive noise in the EEG/EMG signals that interfered with scoring were eliminated from further analyses.

Once we obtained reliable vigilance state classification for each 2 s epoch, the data were then binned into 3 hr time windows and the percent time spent in either REM, NREM, or wake were calculated along with the total number and duration of each bout of these states. The final 3 hr bin (ZT 9:00 thru 11:59) was excluded from analysis because this is when recordings were interrupted for animal husbandry and drug treatments. A “bout” was defined as a contiguous block of 2 s epochs assigned to the same state, and the total number of bouts is the sum of bouts over the 3 hr bin. The bout duration metric was calculated by averaging the number of 2 s epochs per bout, and then multiplying this number by 2 s (i.e., the duration of each 2s epoch).

### Behavioral testing procedures

#### Sucrose preference test

To pre-expose mice to individual housing and the two-bottle choice (sucrose preference test) procedure, mice were anesthetized and ear punched for future identification, then after a short recovery time, were separated into individual cages with a nestlet and food ad lib. Using standard home cage lids equipped for inserting two sippers/bottles side by side, we gave mice two water bottles for the first 23 hr period. Bottles were placed in the lid ∼15-30min before lights off. After the first 23hrs with two water bottles, we provided mice with one bottle containing water and one containing 1% sucrose in facility water. Both bottles were weighed to compare later for obtaining amount of liquids consumed, and the side which contained sucrose and water was counterbalanced for all mice. The mice were then placed in a back room with the same lights on and off schedule and a white noise machine to mask any outside noises. The room was not entered for the duration of drinking for any drinking period. The data for this first session not analyzed because of initial side biases of mice. That is, this first session was an opportunity for mice to sample each liquid and learn that there were two different liquids. The same procedure was performed for two more 23hr periods (i.e., 2 days of pretreatment), with side assignment of sucrose and water bottles reversed each night. After finishing the pretest experiments, mice were again group housed with their original housemates (4 per cage) and were allowed to reacclimate to group housing for 36 hours. Then the mice were given the 6-day THC (10mg/kg) treatment, with 2 mice in each cage receiving THC injections and the other two mice receiving vehicle injections each day. Twenty-four hours after the final injection, the mice were again separated into individual cages and provided two bottle choice of the same sucrose and water solutions, and left in the same room, under the same conditions, for the same time periods as in the pretest. Bottles were measured every 24 hours for the next six days of abstinence. Sucrose preference was inferred by the percent sucrose consumed of total liquid volume consumed.

#### Conditioned operant sucrose seeking

Mice were group housed, four per cage, throughout the experiment. Mice were acclimated to water restriction over several days to potentiate acquisition of operant sucrose seeking. Experiment sessions were conducted in standard operant conditioning chambers (Med Associates, ENV-307W). Each chamber was equipped with two levers, two cue lamps, a house lamp, an audio stimulus generator, a syringe pump for reward delivery, and a reward port equipped with a contact lickometer (ENV-303RMW-4.25). Operant sessions were conducted between ZT700 and ZT1000.

*First training procedure:* Both levers are available throughout the 30-minute session. A lever press on either lever resulted in a two-second auditory cue (CS+), which began after the lever-press, followed by delivery of the sucrose solution (∼20 uL of 8% sucrose, 1% maltodextrin, in facility water) at the end of the cue. Additional lever-presses had no result until mice retrieved the sucrose reward as detected by breaking the photobeams in the sucrose port. Occasionally sucrose solution droplets were added to the levers to facilitate approach to the lever and operant responding. Mice had to successfully earn >21 rewards during the session before moving to the second training procedure.

Second training procedure: This procedure was identical to the first training procedure, except a second auditory cue was added to serve as a predictor of no sucrose reward (CS-). CS+ occurred after 50% of lever presses and CS-after the other 50% in a random fashion. Mice quickly learned to discriminate between CS+ and CS- and after 3 sessions were no longer water restricted.

Third training procedure: This procedure was identical to the second training procedure, but mice had access to water ad-lib in home cage. This lasted an additional 5 daily sessions, with the final two sessions comprising the data for pretreatment condition epoch.

#### Food and water intake and locomotor activity monitoring

Mice were individually housed in Sable Systems International Promethion 3721 Mouse Cage (8.1 × 14.4 × 5.5 in). Cages were equipped with load-cells to monitor food and water intake each second (part # MM-1) and cages were surrounded by photobeams (part # - BXYZ), allowing for precise measurement of consumption and locomotor activity. Cages were lined with 1/8” Teklad corncob bedding (Catalog# 7092). Mice were given a nestlet and an enrichment device (a solid, plastic cube which while placed beneath the food hopper deterred nesting beneath the feeder). After a five-day acclimation period, mice began the treatment and abstinence procedures that were identical to sleep experiments. Mice were also weighed daily at ZT11:00 beginning just prior to vehicle treatment.

#### Bottle brush test

The bottle brush test (BBT) was adapted from Riittinen et al. (1986), Lagerspetz and Portin (1968), and Kimbrough et al. (2017). BBT sessions were conducted between ZT1300 and ZT1500. Mice were handled by the experimenter for 5 minutes per day for 3 days prior to beginning the BBT experiment. On test days, home cages were covered with a sheet to block ambient light and then transferred to the behavioral suite. The behavioral suite was equipped with white-noise sound generators and was illuminated with red-ambient light only. The testing arena was separated from the ‘holding’ area by a door so that mice being tested and cage mates were sensory-isolated from each other. The experimenter wore a red-light headlamp. Ten min before its test session, a mouse was put in a fresh home cage with clean bedding and allowed to habituate. Next, the cage was picked up and transferred to the BBT recording area, which was a roofless, white-walled 2 ft x 2ft x 2ft enclosure, and the walls of the animal’s cage were extended by stacking on top an additional “floorless” identical cage. These two testing area features achieved, respectively, 1) separating the animal visually from the rest of the behavioral room and 2) preventing the mouse from jumping out of its test cage while the BBT was conducted.

Mice were group housed, 4 per cage, for the duration of the experiment except for separation one-by-one as described above during the BBT itself. The test itself was performed in its entirety four separate times. First on the third day of handling (pre-exposure), then again 48hrs later (pretreatment), followed by the 6-day chronic THC/vehicle treatment procedure, and then on days 2 and 6 of abstinence. Treatment groups were assigned randomly, with 2 mice from each cage receiving THC and the other 2 mice vehicle.

During the test, irritability behaviors were elicited by a rotating bottle brush (NSN 7920-00-297-1510) with brown bristles that extended ¾ inch radially for 3.5 inches lengthwise, which was attached to a ¼ inch diameter x 2 ft metal rod that allowed the experimenter to remain at a distance from the test cage. Each attack consisted of five “stages” in the following sequence: 1) rotating brush (180 degrees about 2x/sec; with brush spinning axially via twisting of the rod in the experimenter’s fingers) toward the mouse from the opposite end of the cage, 2) continuing to rotate the brush, but now against the animal’s whiskers, 3) rotating brush moving away from mouse back to opposite end of cage, 4) rotating brush at the starting position, 5) brush at starting position without rotating. Each stage was designed to elicit various aspects of murine irritability responses, and stage lasted 2 seconds, except the 5th stage that was prolonged until the mouse returned to its end of the cage or after 5 seconds. This was repeated for each animal 10 times per test, with 10-15 second intertrial intervals. Cage-mates were habituated in a staggered manner and kept outside the testing room until testing, with the order between treatment groups counterbalanced within and between cages. Bottle brushes were switched between mice to prevent responses to conspecific scents. All brushes were cleaned with ethanol and left to dry before being used for another testing day.

BBT sessions were recorded using Bonsai RX software (10.3389/fninf.2015.00007) to record synchronous video and audio. Video (30 fps) was recorded using two standard USB web cameras without infrared filters (E.L.P.; Ailipu Technology Co., China) mounted above and to the side of the testing cage, and ultrasonic audio (384khz) was recorded from a high-frequency microphone (M500; Pettersson Elektronik AB, Sweden) mounted 18 inches above the testing cage. Behaviors were scored manually via top and side recorded video. Scores for irritability-like behavior were determined by measuring aggressive and defensive behavior. Responses were designated as aggressive or defensive as described by Kimbrough et al. (2017). Aggressive responses scored were smelling, biting, boxing, following, exploring, and tail rattling. Defensive/stress-induced responses were escaping, digging, jumping, climbing, defecating, vocalizing, and grooming.

### Plasma corticosterone collection

Pretreatment samples were collected the day before the initial 10mg/kg THC injection of the treatment paradigm, while early and late samples were collected two days and six days after the final treatment, respectively. Mice were restrained using a Tailveiner Restrainer (TV-150) such that the mouse was placed into the device tail-first. The tail was gently rubbed with a hand warmer for 5 seconds to increase blood flow. A carbon steel scalpel blade (size 11, Integra) was used to snip the lateral tail vein and blood was collected using a heparinized Micro-Hematocrit capillary tube (Fisher Scientific, PA). Pressure was applied to the snip site to stop excessive bleeding and the mouse was placed in a clean new cage within the room where sample collection occurred. Cage mates were kept outside the blood sampling room until blood collection. Sampling took no longer than 2 minutes per mouse. The capillary tubes were spun in a DM1424 Hematocrit Centrifuge (SCILOGEX, LLC, CT, USA) to separate red blood cell and plasma components. 15-30 uL of plasma was transferred into an Eppendorf tube and stored in a -20 C freezer until further analysis.

### Statistical analysis

For sleep, intake/locomotion, sucrose preference test, and operant tasks, the pretreatment, early, and late withdrawal metric were the average of the two pretreatment sessions, first two and last two days of abstinence sessions, respectively. For all other experiments we only recorded metrics during one day for pretreatment, second and sixth day of abstinence, so analysis reflects those single sessions.

For sleep studies we analyzed the data by fitting a mixed model as implemented in GraphPad Prism 9.0. This mixed model uses a compound symmetry covariance matrix and is fit using Restricted Maximum Likelihood (REML). In the absence of missing values, this method gives the same P values and multiple comparisons tests as repeated measures ANOVA. In the presence of missing values, the results can be interpreted like a repeated measures ANOVA. For these data a three-way (2_photoperiod_ x 2_epoch_ x 2_treatment_) repeated measures mixed-effects model was used, allowing us to analyze the data set despite missing values for some subjects on some recording sessions. These occurred at random due to scenarios such as, mice breaking recording tethers, moisture accumulation from water during a recording session leading to static electricity artifacts/noisy recordings, and headcaps or recording tethers becoming detached. For consistency, the same analysis was used for 24hr intake and locomotion behavior recordings, and all other behavioral studies used a two-way 2_epoch_ x 2_treatment_ repeated measures mixed-effects model. When significant (p<0.05) main and interaction effects relating to treatment were detected, we followed up with Holm-Šídák’s multiple comparisons post-hoc test to compare means of metrics differing by only one factor. Since photoperiod has clear effects on sleep regardless of treatments or epochs, we only report significant post-hoc tests comparing treatment effects and within-subjects effects related to epoch. For voltammetry experiments a two-way ANOVA were used for statistical comparisons between groups.

---

**Supplemental Figure 1.**
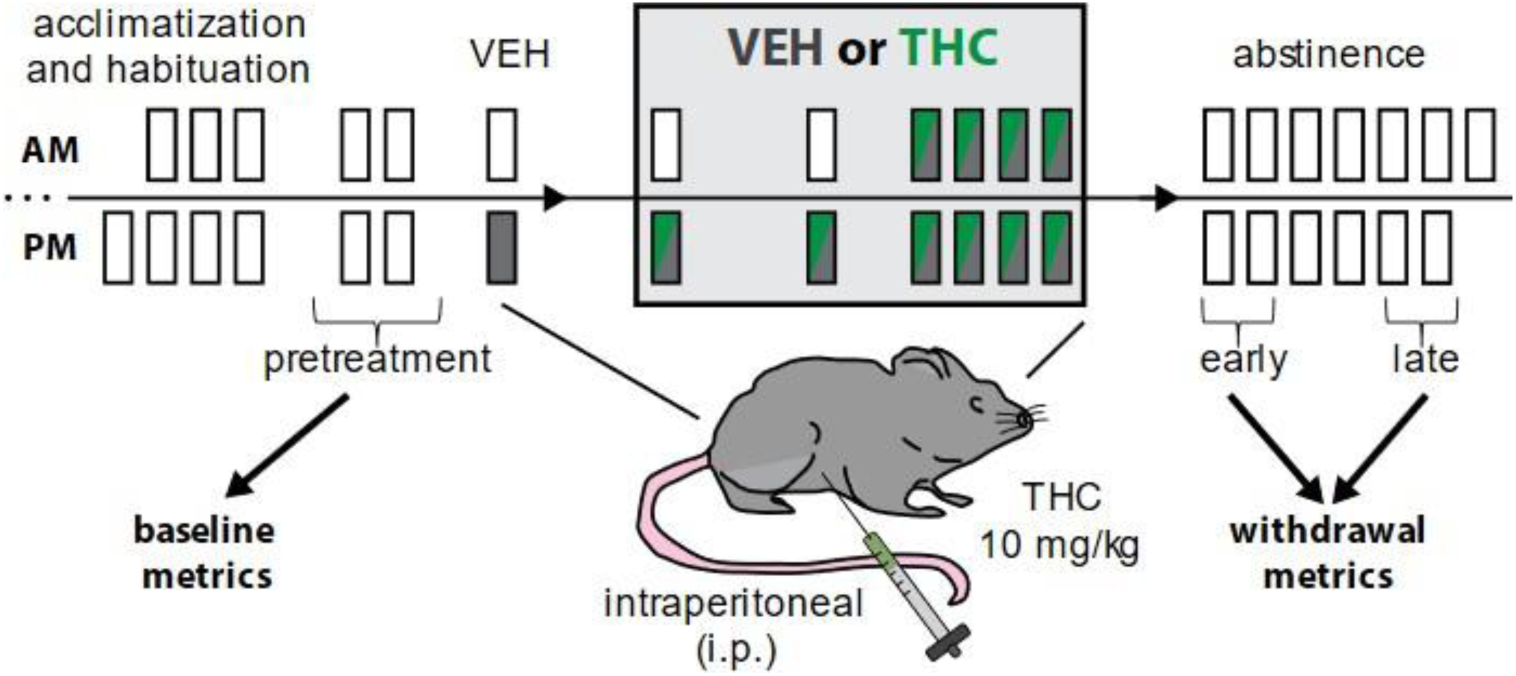
General timeline used to model chronic cannabis/THC use in mice. Time points allow comparison of pretreatment epochs to early and late withdrawal, and initial vehicle injection to ‘first’ and ‘last’ THC or VEH injections during treatment epoch.

**Supplemental Figure 2.**
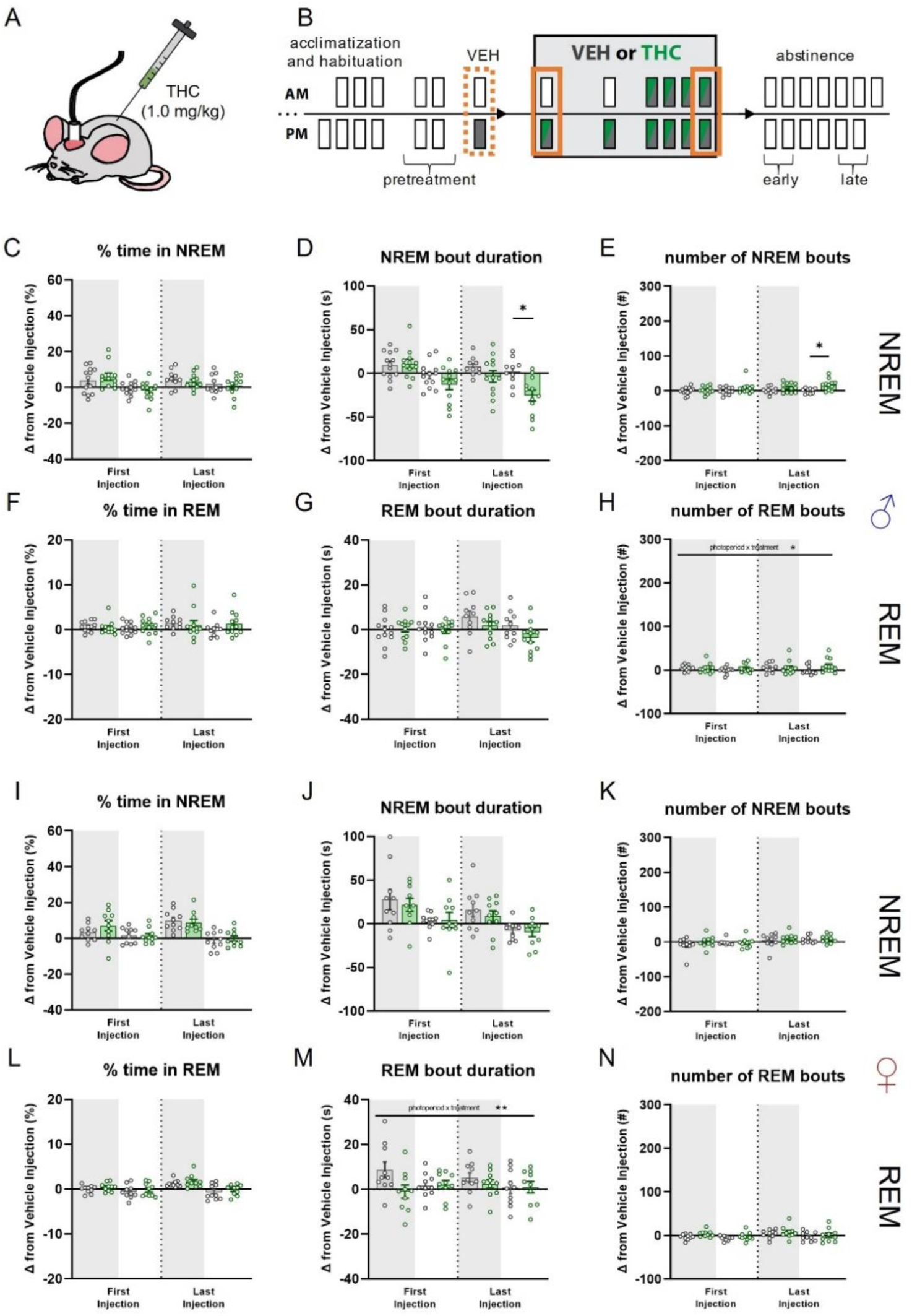
Effects of acute and chronic low dose THC administration on sleep in male and female mice. (A) Cartoon illustrating i.p. administration of lower dose, 1.0mg/kg THC. (B) Timeline illustrating treatment regimen and epochs contributing to analysis. Dotted orange line indicates vehicle treatment epoch that acute and chronic administration epochs are compared. (C) Comparison between effects of THC or VEH on changes from vehicle injection in percent time spent in NREM sleep in male mice. No significant effects were detected. (D) Comparison between effects of THC or VEH on changes from vehicle injection in NREM bout duration in male mice. There is a significant treatment effect (F_treatment_(1,23) = 5.834; p = 0.0241), with posthoc tests finding differences between-groups during Last Injection-LON (p* = 0.0179). (E) Comparison between effects of THC or VEH on changes from vehicle injection in number of NREM bouts in male mice. There is a significant treatment effect (F_treatment_(1,23) = 7.398; p = 0.0122), with posthoc tests finding differences between-groups during Last Injection-LON (p* = 0.0428). (F) - (H) Same metrics as (C-D) but for REM sleep in male mice. There were no significant effects detected for percent time in REM (F) or REM bout duration (G). A significant effect of treatment was detected (F_treatment_(1,22) = 9.312; p = 0.0059), but posthoc tests did not detect between or within group effects. (I) - (N) Same metrics as (C)-(H) but in female mice. A significant interaction for REM bout duration was detected (F_photoperiod x treatment_(1,18) = 7.939; p = 0.0114), but posthoc tests did not detect between or within group effects.

**Supplemental Figure 3.**
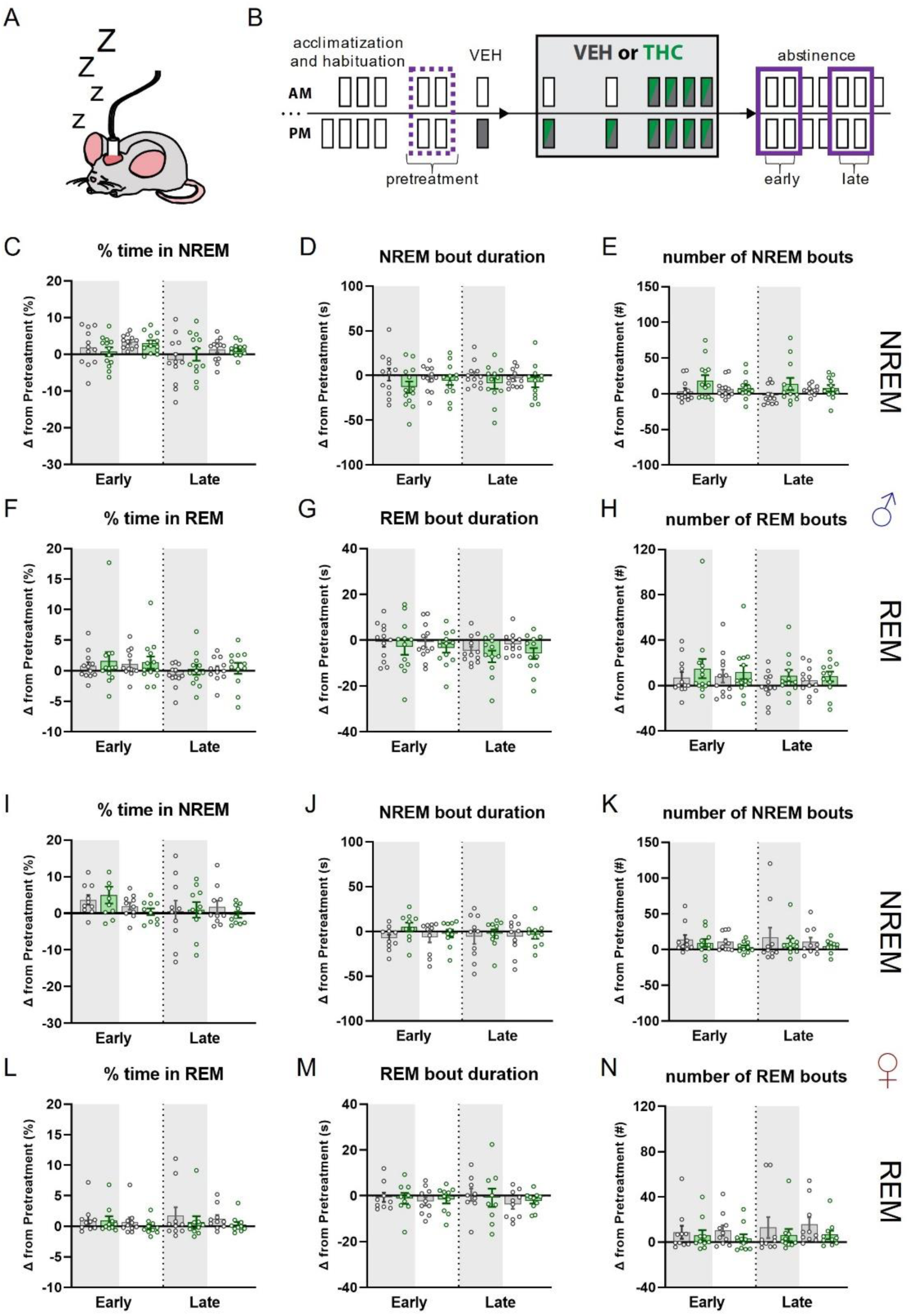
Effects of chronic low dose (1.0 mg/kg) THC administration on sleep in male and female mice during early and late abstinence. (A) Cartoon illustrating chronically tethered mouse during epochs where no injection is given, i.e., pretreatment and abstinence. (B) Timeline illustrating treatment regimen and epochs contributing to analysis. Dotted purple line indicates average of two pretreatment epochs that average of days 1 and 2 of early and days 5 and 6 of late abstinence are compared. (C) - (N) No significant effects were detected in either.

**Supplemental Figure 4.**
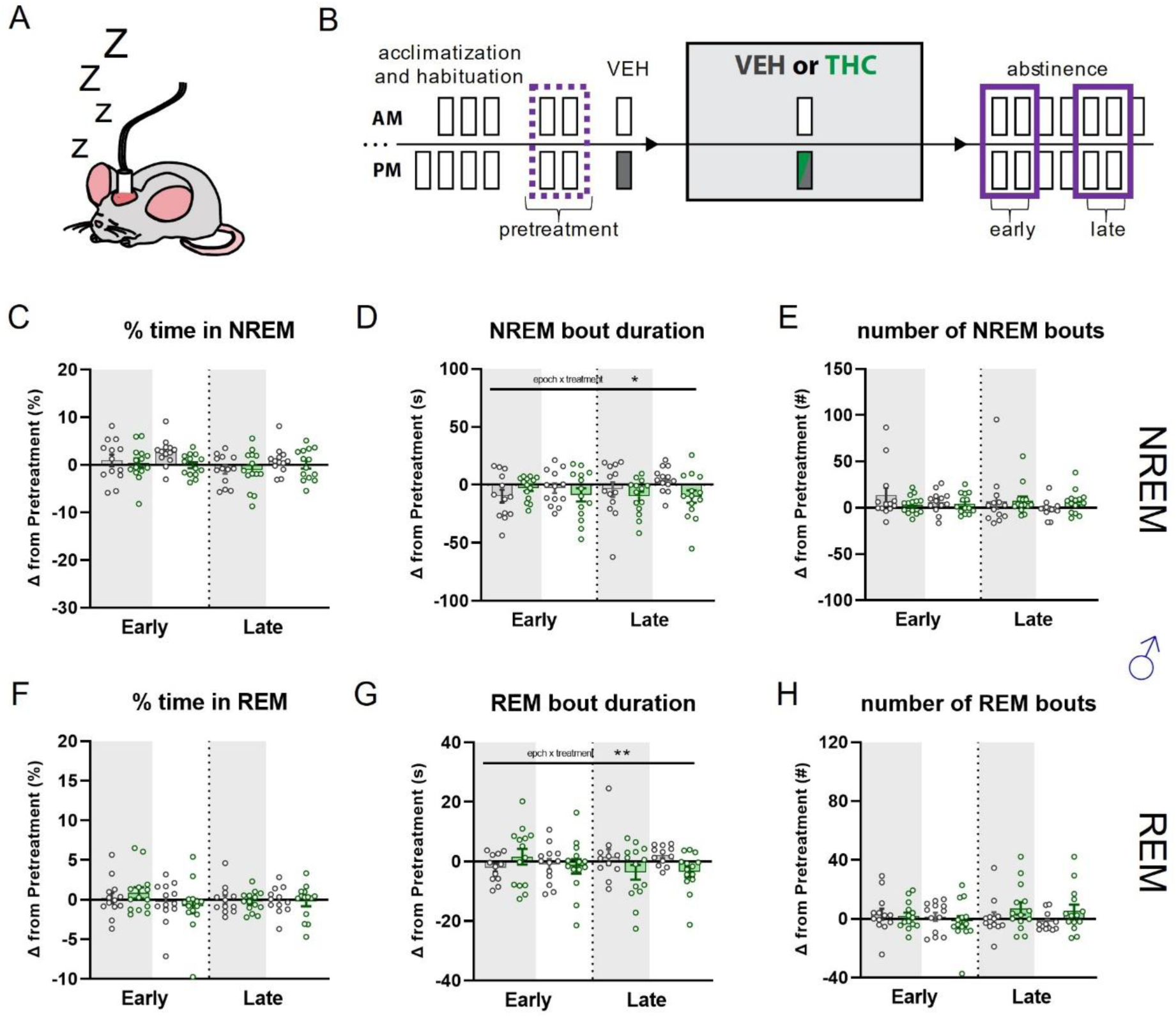
Effects acute THC administration on sleep in male mice during early and late abstinence. (A) Cartoon illustrating chronically tethered mouse during epochs where no injection is given, i.e., pretreatment and abstinence. (B) Timeline illustrating treatment regimen and epochs contributing to analysis. Dotted purple line indicates average of two pretreatment epochs that average of days 1 and 2 of early and days 5 and 6 of late abstinence are compared. (C) - (H) NREM and REM metrics. Significant interaction effects were detected for bout duration in NREM (D; F_epoch x treatment_(1,25) = 5.388; p = 0.0287) and REM (G; F_epoch x treatment_(1,25) = 7.828; p = 0.0098). Posthoc tests for these metrics did not reveal any significant between or within group differences.

**Supplemental Figure 5.**
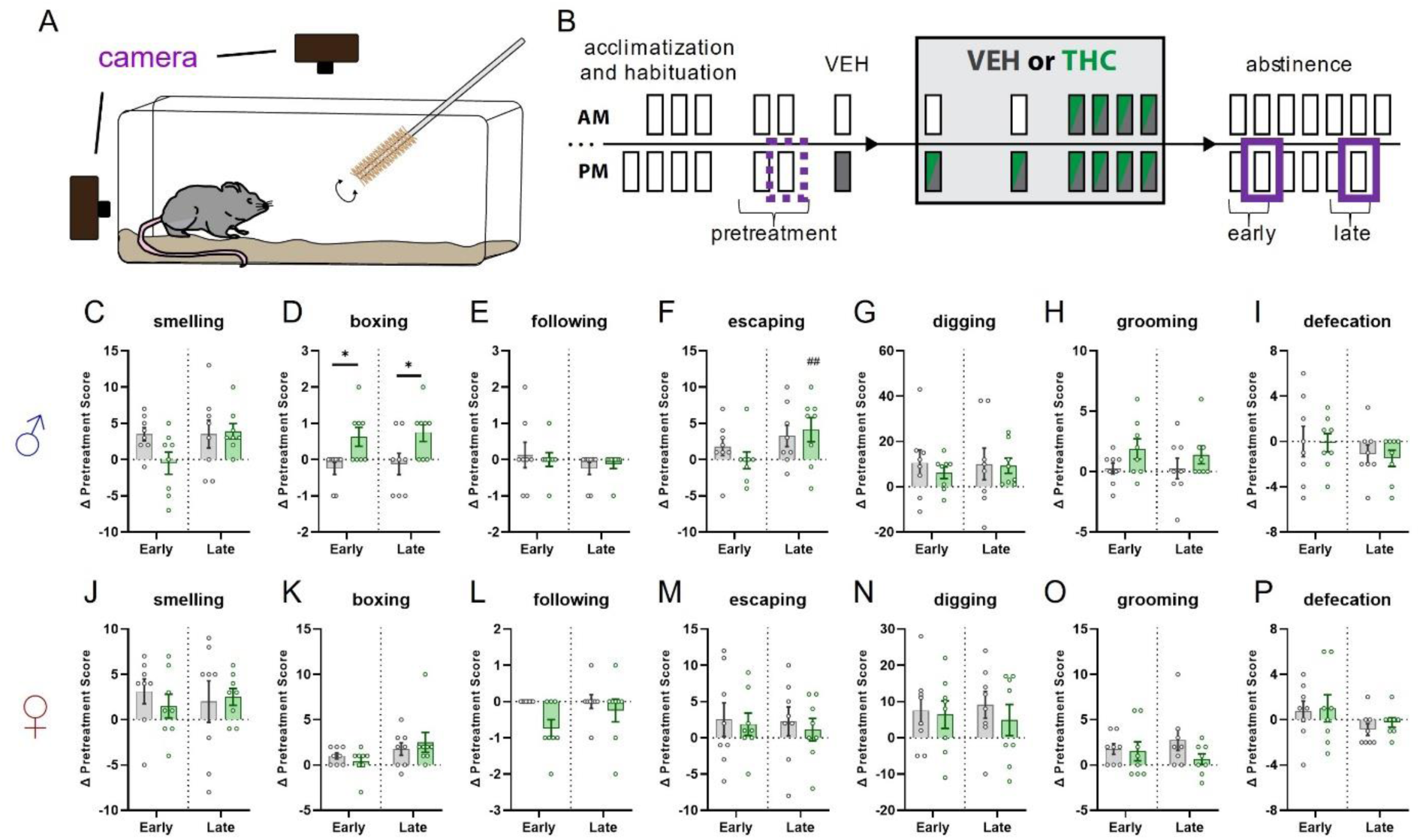
Effects of chronic THC administration on irritability-related behaviors during Early and Late abstinence in male and female mice. (A) Cartoon illustrating bottle brush test procedure for measuring irritability-related responses. (B) Timeline illustrating treatment regimen and epochs contributing to analysis. Dotted purple line indicates pretreatment session that Early and Late abstinence sessions are compared. (C) - (I) Change in irritability-related behavioral metrics during Early and Late abstinence in male mice. Significant effect of treatment was detected for boxing behavior (D; F_treatment_(1,14) = 12.47; p = 0.0015) with post-hoc tests finding significant group differences in Early and Late abstinence (p* = 0.0187). Significant effect of epoch was detected for escaping behavior (F; F_epoch_(1,14) = 8.211; p = 0.0125) with post-hoc tests finding within group difference in THC treated mice (p^##^ = 0.0096). (J) - (P) Same metrics as above but for female mice. No significant effects were detected.

**Supplemental Figure 6.**
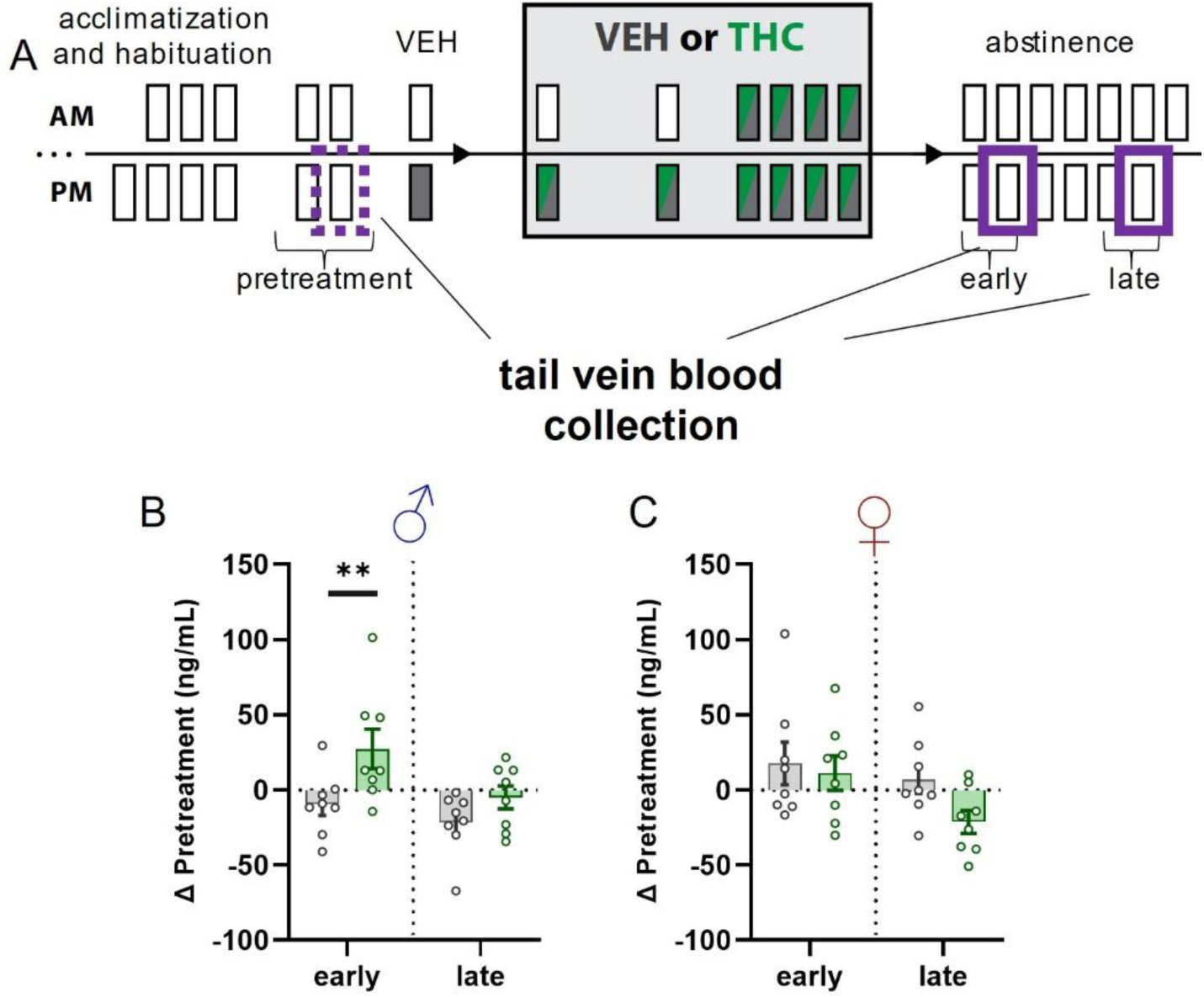
Effect of chronic THC treatment on circulating plasma corticosterone levels in male in female mice during Early and Late abstinence. (A) Timeline illustrating treatment regimen and epochs contributing to analysis. Dotted purple line indicates pretreatment sample that Early and Late abstinence samples are compared. (B) Change in plasma CORT from pretreatment levels in Early and Late abstinence in male mice. A significant effect of treatment was detected F_treatment_(1,14) = 6.115; p = 0.0332) with posthoc tests finding between-group differences in Early abstinence (p** = 0.0168). (C) Same as (B) but for female mice. No significant effects were detected.

## Notes

### Competing Interest Statement

The authors have declared no competing interest.

## REFERENCES

1. United Nations Office on Drugs and Crime, World Drug Report 2019, U.N.N. York, Editor. 2019, United Nations New York: New York.

2. Smart, R. and R.L. Pacula, Early evidence of the impact of cannabis legalization on cannabis use, cannabis use disorder, and the use of other substances: Findings from state policy evaluations. Am J Drug Alcohol Abuse, 2019. 45(6): p. 644–663.

3. Kesner, A.J. and D.M. Lovinger, Cannabis use, abuse, and withdrawal: Cannabinergic mechanisms, clinical, and preclinical findings. J Neurochem, 2021. 157(5): p. 1674–1696.

4. Jones, R.T., N. Benowitz, and J. Bachman, Clinical studies of cannabis tolerance and dependence. Ann N Y Acad Sci, 1976. 282: p. 221–39.

5. Compton, D.R., W.L. Dewey, and B.R. Martin, Cannabis dependence and tolerance production. Adv Alcohol Subst Abuse, 1990. 9(1-2): p. 129–47.

6. American Psychiatric Association, Diagnostic and statistical manual of mental disorders (5th ed.). 2013, Washington, DC.

7. Huestis, M.A., et al., Blockade of effects of smoked marijuana by the CB1-selective cannabinoid receptor antagonist SR141716. Arch Gen Psychiatry, 2001. 58(4): p. 322–8.

8. Devane, W.A., et al., Determination and characterization of a cannabinoid receptor in rat brain. Mol Pharmacol, 1988. 34(5): p. 605–13.

9. Lichtman, A.H., et al., Opioid and cannabinoid modulation of precipitated withdrawal in delta(9)-tetrahydrocannabinol and morphine-dependent mice. J Pharmacol Exp Ther, 2001. 298(3): p. 1007–14.

10. Ledent, C., et al., Unresponsiveness to cannabinoids and reduced addictive effects of opiates in CB1 receptor knockout mice. Science, 1999. 283(5400): p. 401–4.

11. Martellotta, M.C., et al., Self-administration of the cannabinoid receptor agonist WIN 55,212-2 in drug-naive mice. Neuroscience, 1998. 85(2): p. 327–30.

12. Haney, M., et al., Abstinence symptoms following oral THC administration to humans. Psychopharmacology (Berl), 1999. 141(4): p. 385–94.

13. Haney, M., et al., Abstinence symptoms following smoked marijuana in humans. Psychopharmacology (Berl), 1999. 141(4): p. 395–404.

14. Vandrey, R., et al., Cannabis withdrawal in adolescent treatment seekers. Drug Alcohol Depend, 2005. 78(2): p. 205–10.

15. Vandrey, R.G., et al., A within-subject comparison of withdrawal symptoms during abstinence from cannabis, tobacco, and both substances. Drug Alcohol Depend, 2008. 92(1-3): p. 48–54.

16. Budney, A.J., et al., Comparison of cannabis and tobacco withdrawal: severity and contribution to relapse. J Subst Abuse Treat, 2008. 35(4): p. 362–8.

17. Budney, A.J., et al., The time course and significance of cannabis withdrawal. J Abnorm Psychol, 2003. 112(3): p. 393–402.

18. Budney, A.J., et al., Oral delta-9-tetrahydrocannabinol suppresses cannabis withdrawal symptoms. Drug Alcohol Depend, 2007. 86(1): p. 22–9.

19. Kouri, E.M. and H.G. Pope, Jr., Abstinence symptoms during withdrawal from chronic marijuana use. Exp Clin Psychopharmacol, 2000. 8(4): p. 483–92.

20. Kouri, E.M., H.G. Pope, Jr., and S.E. Lukas, Changes in aggressive behavior during withdrawal from long-term marijuana use. Psychopharmacology (Berl), 1999. 143(3): p. 302–8.

21. Budney, A.J., P.L. Novy, and J.R. Hughes, Marijuana withdrawal among adults seeking treatment for marijuana dependence. Addiction, 1999. 94(9): p. 1311–22.

22. Budney, A.J., et al., Marijuana abstinence effects in marijuana smokers maintained in their home environment. Arch Gen Psychiatry, 2001. 58(10): p. 917–24.

23. Crowley, T.J., et al., Cannabis dependence, withdrawal, and reinforcing effects among adolescents with conduct symptoms and substance use disorders. Drug and Alcohol Dependence, 1998. 50(1): p. 27–37.

24. Coughlin, L.N., et al., Progression of cannabis withdrawal symptoms in people using medical cannabis for chronic pain. Addiction, 2021. 116(8): p. 2067–2075.

25. Bahji, A., et al., Prevalence of Cannabis Withdrawal Symptoms Among People With Regular or Dependent Use of Cannabinoids: A Systematic Review and Meta-analysis. JAMA Netw Open, 2020. 3(4): p. e202370.

26. Babson, K.A., M.T. Boden, and M.O. Bonn-Miller, The impact of perceived sleep quality and sleep efficiency/duration on cannabis use during a self-guided quit attempt. Addict Behav, 2013. 38(11): p. 2707–13.

27. Babson, K.A., et al., Poor sleep quality as a risk factor for lapse following a cannabis quit attempt. J Subst Abuse Treat, 2013. 44(4): p. 438–43.

28. Bonn-Miller, M.O., et al., Priority Considerations for Medicinal Cannabis-Related Research. Cannabis Cannabinoid Res, 2019. 4(3): p. 139–157.

29. Gorelick, D.A., et al., Around-the-clock oral THC effects on sleep in male chronic daily cannabis smokers. Am J Addict, 2013. 22(5): p. 510–4.

30. Hosko, M.J., M.S. Kochar, and R.I. Wang, Effects of orally administered delta-9-tetrahydrocannabinol in man. Clin Pharmacol Ther, 1973. 14(3): p. 344–52.

31. Freemon, F.R., The effect of δ 9-tetrahydrocannabinol on sleep. Psychopharmacologia, 1974. 35: p. 39–44.

32. Cousens, K. and A. DiMascio, (-) Delta 9 THC as an hypnotic. An experimental study of three dose levels. Psychopharmacologia, 1973. 33(4): p. 355–64.

33. Pava, M.J., A. Makriyannis, and D.M. Lovinger, Endocannabinoid Signaling Regulates Sleep Stability. PLoS One, 2016. 11(3): p. e0152473.

34. Freemon, F.R., The Effect of Chronically Administered Delta-9-Tetrahydrocannabinol Upon the Polygraphically Monitored Sleep of Normal Volunteers. Drug and Alcohol Dependence, 1982. 10(4): p. 345–353.

35. Bolla, K.I., et al., Polysomnogram changes in marijuana users who report sleep disturbances during prior abstinence. Sleep Med, 2010. 11(9): p. 882–9.

36. Vandrey, R., et al., Sleep disturbance and the effects of extended-release zolpidem during cannabis withdrawal. Drug Alcohol Depend, 2011. 117(1): p. 38–44.

37. Aceto, M.D., et al., Cannabinoid precipitated withdrawal by the selective cannabinoid receptor antagonist, SR 141716A. Eur J Pharmacol, 1995. 282(1-3): p. R1–2.

38. Maldonado, R. and F. Rodriguez de Fonseca, Cannabinoid addiction: behavioral models and neural correlates. J Neurosci, 2002. 22(9): p. 3326–31.

39. Lichtman, A.H. and B.R. Martin, Marijuana withdrawal syndrome in the animal model. J Clin Pharmacol, 2002. 42(S1): p. 20S–27S.

40. Gonzalez, S., M. Cebeira, and J. Fernandez-Ruiz, Cannabinoid tolerance and dependence: a review of studies in laboratory animals. Pharmacol Biochem Behav, 2005. 81(2): p. 300–18.

41. Trexler, K.R., et al., Novel behavioral assays of spontaneous and precipitated THC withdrawal in mice. Drug Alcohol Depend, 2018. 191: p. 14–24.

42. Volkow, N.D. and M. Morales, The Brain on Drugs: From Reward to Addiction. Cell, 2015. 162(4): p. 712–25.

43. Wise, R.A., Dopamine, learning and motivation. Nat Rev Neurosci, 2004. 5(6): p. 483–94.

44. Di Chiara, G., The role of dopamine in drug abuse viewed from the perspective of its role in motivation. Drug Alcohol Depend, 1995. 38(2): p. 95–137.

45. Volkow, N.D., J.S. Fowler, and G.J. Wang, Role of dopamine in drug reinforcement and addiction in humans: results from imaging studies. Behav Pharmacol, 2002. 13(5-6): p. 355–66.

46. Alonso, I.P., et al., Dopamine transporter function fluctuates across sleep/wake state: potential impact for addiction. Neuropsychopharmacology, 2020.

47. Eban-Rothschild, A., et al., VTA dopaminergic neurons regulate ethologically relevant sleep-wake behaviors. Nat Neurosci, 2016.

48. Dong, H., et al., Dorsal Striatum Dopamine Levels Fluctuate Across the Sleep-Wake Cycle and Respond to Salient Stimuli in Mice. Front Neurosci, 2019. 13: p. 242.

49. Covey, D.P., et al., Endocannabinoid modulation of dopamine neurotransmission. Neuropharmacology, 2017. 124: p. 52–61.

50. Sami, M.B., E.A. Rabiner, and S. Bhattacharyya, Does cannabis affect dopaminergic signaling in the human brain? A systematic review of evidence to date. Eur Neuropsychopharmacol, 2015. 25(8): p. 1201–24.

51. Bloomfield, M.A., et al., The effects of Delta(9)-tetrahydrocannabinol on the dopamine system. Nature, 2016. 539(7629): p. 369–377.

52. Koob, G.F. and N.D. Volkow, Neurocircuitry of addiction. Neuropsychopharmacology, 2010. 35(1): p. 217–38.

53. Hasegawa, H., et al., The subcortical belly of sleep: New possibilities in neuromodulation of basal ganglia? Sleep Med Rev, 2020. 52: p. 101317.

54. Bass, C.E. and B.R. Martin, Time course for the induction and maintenance of tolerance to Delta(9)-tetrahydrocannabinol in mice. Drug Alcohol Depend, 2000. 60(2): p. 113–9.

55. Cooper, Z.D. and R.M. Craft, Sex-Dependent Effects of Cannabis and Cannabinoids: A Translational Perspective. Neuropsychopharmacology, 2018. 43(1): p. 34–51.

56. Abrahao, K.P., M.J. Pava, and D.M. Lovinger, Dose-dependent alcohol effects on electroencephalogram: Sedation/anesthesia is qualitatively distinct from sleep. Neuropharmacology, 2020. 164: p. 107913.

57. Bolla, K.I., et al., Sleep disturbance in heavy marijuana users. Sleep, 2008. 31(6): p. 901–8.

58. Babson, K.A. and M.O. Bonn-Miller, Sleep Disturbances: Implications for Cannabis Use, Cannabis Use Cessation, and Cannabis Use Treatment. Current Addiction Reports, 2014. 1(2): p. 109–114.

59. Schlienz, N.J. and R. Vandrey, Cannabis Withdrawal, in Cannabis Use Disorders. 2019. p. 93–102.

60. Lagerspetz, K. and R. Portin, Simulation of cues eliciting aggressive responses in mice at two age levels. J Genet Psychol, 1968. 113(1st Half): p. 53–63.

61. Riittinen, M.L., et al., Impoverished rearing conditions increase stress-induced irritability in mice. Dev Psychobiol, 1986. 19(2): p. 105–11.

62. Pawlyk, A.C., et al., Stress-induced changes in sleep in rodents: models and mechanisms. Neurosci Biobehav Rev, 2008. 32(1): p. 99–117.

63. Sadler, A.M. and S.J. Bailey, Repeated daily restraint stress induces adaptive behavioural changes in both adult and juvenile mice. Physiol Behav, 2016. 167: p. 313–323.

64. Manza, P., D. Tomasi, and N.D. Volkow, Subcortical Local Functional Hyperconnectivity in Cannabis Dependence. Biol Psychiatry Cogn Neurosci Neuroimaging, 2018. 3(3): p. 285–293.

65. Volkow, N.D., et al., Decreased dopamine brain reactivity in marijuana abusers is associated with negative emotionality and addiction severity. Proc Natl Acad Sci U S A, 2014. 111(30): p. E3149–56.

66. Nicola, S.M., F.W. Hopf, and G.O. Hjelmstad, Contrast enhancement: a physiological effect of striatal dopamine? Cell Tissue Res, 2004. 318(1): p. 93–106.

67. Diana, M., et al., Mesolimbic dopaminergic decline after cannabinoid withdrawal. Proc Natl Acad Sci U S A, 1998. 95(17): p. 10269–73.

68. Mohebi, A., et al., Dissociable dopamine dynamics for learning and motivation. Nature, 2019. 570(7759): p. 65–70.

69. Mateo, Y., et al., Endocannabinoid Actions on Cortical Terminals Orchestrate Local Modulation of Dopamine Release in the Nucleus Accumbens. Neuron, 2017. 96(5): p. 1112–1126 e5.

70. Cachope, R. and J.F. Cheer, Local control of striatal dopamine release. Front Behav Neurosci, 2014. 8: p. 188.

71. Patriarchi, T., et al., An expanded palette of dopamine sensors for multiplex imaging in vivo. Nat Methods, 2020.

72. Bonn-Miller, M.O., K.A. Babson, and R. Vandrey, Using cannabis to help you sleep: heightened frequency of medical cannabis use among those with PTSD. Drug Alcohol Depend, 2014. 136: p. 162–5.

73. Babson, K.A., J. Sottile, and D. Morabito, Cannabis, Cannabinoids, and Sleep: a Review of the Literature. Curr Psychiatry Rep, 2017. 19(4): p. 23.

74. Levin, K.H., et al., Cannabis withdrawal symptoms in non-treatment-seeking adult cannabis smokers. Drug Alcohol Depend, 2010. 111(1-2): p. 120–7.

75. Copersino, M.L., et al., Cannabis withdrawal among non-treatment-seeking adult cannabis users. Am J Addict, 2006. 15(1): p. 8–14.

76. Mondino, A., et al., Acute effect of vaporized Cannabis on sleep and electrocortical activity. Pharmacol Biochem Behav, 2019. 179: p. 113–123.

77. Willinsky, M.D., A. Scotti de Carolis, and V.G. Longo, EEG and behavioral effects of natural, synthetic and biosynthetic cannabinoids. Psychopharmacologia, 1973. 31(4): p. 365–74.

78. Wallach, M.B. and S. Gershon, The effects of Δ8-THC on the EEG, reticular multiple unit activity and sleep of cats. European Journal of Pharmacology, 1973. 24(2): p. 172–178.

79. Barratt, E.S., W. Beaver, and R. White, The effects of marijuana on human sleep patterns. Biol Psychiatry, 1974. 8(1): p. 47–54.

80. Feinberg, I., et al., Effects of marijuana extract and tetrahydrocannabinol on electroencephalographic sleep patterns. Clin Pharmacol Ther, 1976. 19(6): p. 782–94.

81. Pivik, R.T., et al., Delta-9-tetrahydrocannabinol and synhexl: effects on human sleep patterns. Clin Pharmacol Ther, 1972. 13(3): p. 426–35.

82. Feinberg, I., et al., Effects of high dosage delta-9-tetrahydrocannabinol on sleep patterns in man. Clin Pharmacol Ther, 1975. 17(4): p. 458–66.

83. Nicholson, A.N., et al., Effect of Delta-9-tetrahydrocannabinol and cannabidiol on nocturnal sleep and early-morning behavior in young adults. J Clin Psychopharmacol, 2004. 24(3): p. 305–13.

84. Barratt, E.S. and P.M. Adams, Effect of chronic marijuana administration of stages of primate sleep-wakefulness. Biol Psychiatry, 1975. 10(3): p. 315–322.

85. Barratt, E.S. and P.M. Adams, Chronic marijuana usage and sleep-wakefulness cycles in cats. Biol Psychiatry, 1973. 6(3): p. 207–14.

86. Masur, J. and N. Khazan, Induction by Cannabis sativa (marihuana) of rhythmic spike discharges overriding REM sleep electrocorticogram in the rat. Life Sci I, 1970. 9(22): p. 1275–80.

87. Halikas, J.A., et al., A longitudinal study of marijuana effects. Int J Addict, 1985. 20(5): p. 701–11.

88. Karacan, I., et al., Sleep electroencephalographic-electrooculographic characteristics of chronic marijuana users: part I. Ann N Y Acad Sci, 1976. 282: p. 348–74.

89. Pranikoff, K., et al., Effects of marijuana smoking on the sleep EEG. Preliminary studies. JFMA, 1973. 60(3): p. 28–31.

90. Herning, R.I., et al., EEG deficits in chronic marijuana abusers during monitored abstinence: preliminary findings. Ann N Y Acad Sci, 2003. 993: p. 75–8; discussion 79-81.

91. Herning, R.I., W. Better, and J.L. Cadet, EEG of chronic marijuana users during abstinence: Relationship to years of marijuana use, cerebral blood flow and thyroid function. Clinical Neurophysiology, 2008. 119(2): p. 321–331.

92. Herrmann, E.S., E.M. Weerts, and R. Vandrey, Sex differences in cannabis withdrawal symptoms among treatment-seeking cannabis users. Exp Clin Psychopharmacol, 2015. 23(6): p. 415–21.

93. Smith, N.T., A review of the published literature into cannabis withdrawal symptoms in human users. Addiction, 2002. 97(6): p. 621–32.

## Supplemental References

1. Pava, M.J., A. Makriyannis, and D.M. Lovinger, Endocannabinoid Signaling Regulates Sleep Stability. PLoS One, 2016. 11(3): p. e0152473.

2. Abrahao, K.P., M.J. Pava, and D.M. Lovinger, Dose-dependent alcohol effects on electroencephalogram: Sedation/anesthesia is qualitatively distinct from sleep. Neuropharmacology, 2020. 164: p. 107913.

3. Gervasoni, D., et al., Global forebrain dynamics predict rat behavioral states and their transitions. J Neurosci, 2004. 24(49): p. 11137–47.

4. Bastianini, S., et al., SCOPRISM: a new algorithm for automatic sleep scoring in mice. J Neurosci Methods, 2014. 235: p. 277–84.

5. Diniz Behn, C.G., et al., Abnormal sleep/wake dynamics in orexin knockout mice. Sleep, 2010. 33(3): p. 297–306.

6. Weber, F., et al., Control of REM sleep by ventral medulla GABAergic neurons. Nature, 2015. 526(7573): p. 435–8.

